# The comprehensive roadmaps of reprogramming and transformation unveiled antagonistic roles for bHLH transcription factors in the control of cellular plasticity

**DOI:** 10.1101/2020.12.28.424606

**Authors:** A. Huyghe, G. Furlan, J. Schroeder, J. Stüder, F. Mugnier, L. De Matteo, J. Wang, Y. Yu, N. Rama, B. Gibert, P. Wajda, I. Goddard, N. Gadot, M. Brevet, M. Siouda, P. Mulligan, R. Dante, P. Liu, H. Gronemeyer, M. Mendoza-Parra, J. Polo, F. Lavial

## Abstract

Coordinated changes of cellular identity and plasticity are critical for pluripotent reprogramming (PR) and malignant transformation (MT). However, the molecular circuitries orchestrating these modifications, as well as their degree of analogy during reprogramming and transformation, remain unknown. To address this question, we generated “repro-transformable” mice models and dissected comparatively the early events underpinning PR - mediated by Oct4, Sox2, Klf4, c-Myc - and MT - triggered by oncogenic Ras and c-Myc. Transcriptomic analyses allowed the identification of a unique set of markers - the cell surface glycoprotein Thy1 and the transcription factor (TF) Bcl11b - that are commonly downregulated during PR and MT and delineate cellular intermediates (CI) highly amenable to generate pluripotent or malignant derivatives. Comprehensive transcriptomic, epigenomic and functional analyses of different CI, prone or refractory to PR/MT, unveiled that cellular plasticity acquisition precedes the broad extinction of cellular identity. It also demonstrated the existence of specific and shared molecular features of PR and MT while ensuring the identification of broad-range regulators of cellular plasticity. As a proof-of-concept, we revealed that the basic helix-loop-helix (bHLH) class A TF Atoh8 constrains rodent and human iPS cells generation as well as MT and direct neuron conversion. Mechanistically, this TF hampers the reactivation of the pluripotent network during PR and limits the acquisition of phenotypic plasticity during MT. Furthermore, an integrated analysis of Atoh8 genome-wide binding, alongside the other bHLH TFs c-Myc, Ascl1 and MyoD promoting reprogramming/transdifferentiation, unveiled how Atoh8 constrains cellular plasticity by occupying a specific subset of MEF enhancers and by finetuning WNT signalling activity. Collectively, by deconvoluting the early steps of the reprogramming and transformation roadmaps, this integrated study uncoupled changes of cellular plasticity and identity to shed light on novel insights into reprogramming and cancer biology.

**Graphical abstract:** 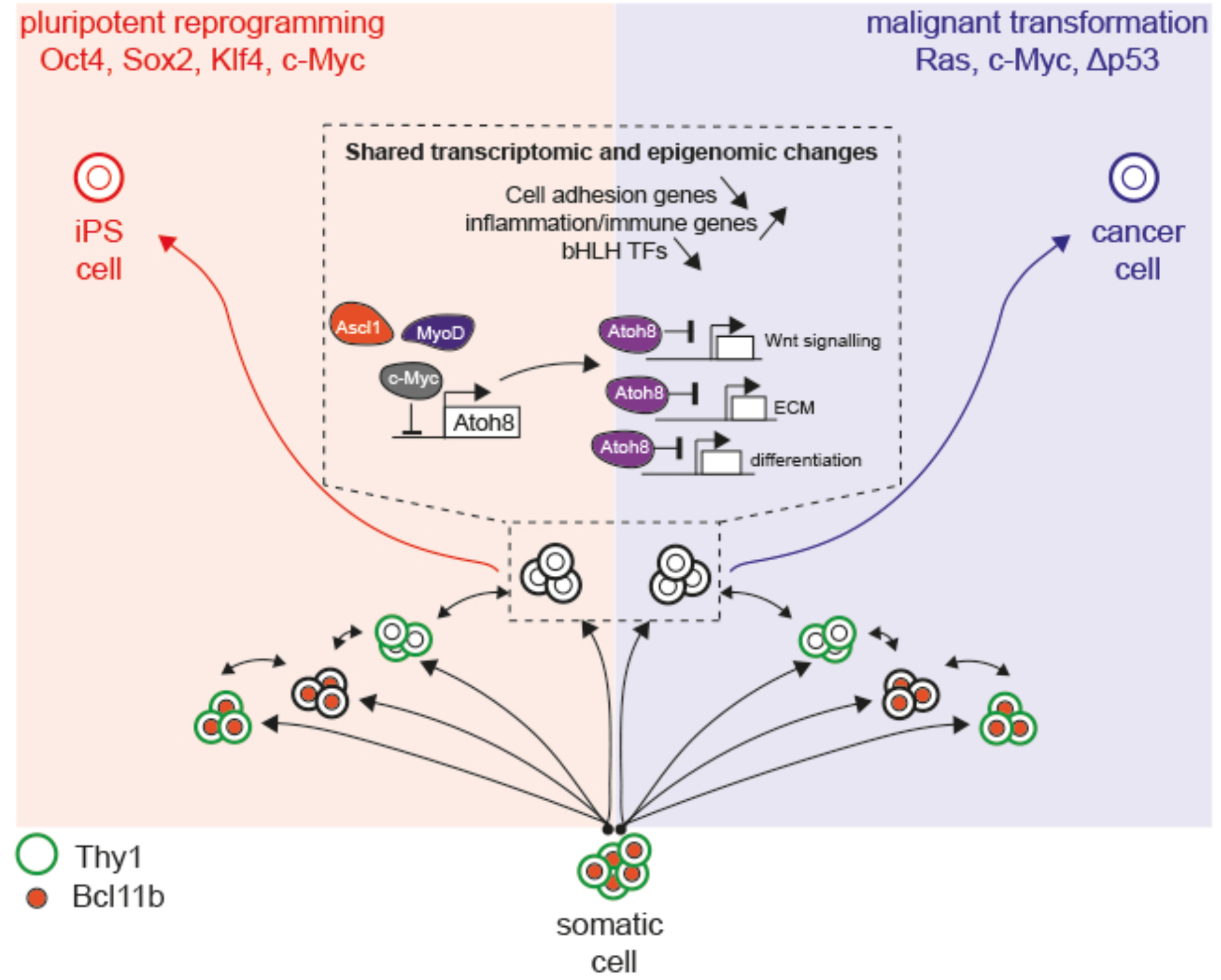

**One-sentence summary:** Comparative roadmaps of cellular plasticity acquisition during pluripotent reprogramming and malignant transformation.

## Introduction

During development, cells within multicellular organisms progressively differentiate into functionally and phenotypically distinct fates. These cellular identities, established by cell typespecific gene expression programs, are remarkably stable and can be sustained over many cell divisions throughout an organism’s lifespan. However, this view of cellular identity as a permanent fixed state has been extensively challenged by the discovery of pluripotent reprogramming (PR). In their seminal report, Takahashi and Yamanaka demonstrated that differentiated cells can be fully converted to pluripotency by a defined set of transcription factors (TFs) (Oct4, Sox2, Klf4, and c-Myc; thereafter named OSKM) (*1*). Mechanistically, OSKM trigger an early and widespread reconfiguration of chromatin states and TF occupancy to orchestrate loss of cellular identity and gain of cellular plasticity in mouse embryonic fibroblasts (MEF), while gradual activation of the pluripotent transcriptional network is observed later during the process of induced pluripotent stem (iPS) cell generation (*2–7*). Despite the definition of the reprogramming roadmaps of diverse human and mouse cells of origin (*8–11*), the molecular mechanisms controlling the stepwise loss of identity and gain of plasticity are still largely unknown, yet they are critical for the coordinated acquisition of induced pluripotency.

Malignant transformation (MT) share some features with PR – both processes are intrinsically constrained by oncogenic barriers, such as cell death and senescence, and are considered to be stochastic and subjected to significant latencies (*12–16*). Importantly, loss of cellular identity and gain of cellular plasticity also emerged as critical steps of MT, as we and others recently reviewed (*16–18*). Indeed, cancer formation frequently relies on the activation of developmental programs that increase cellular plasticity and trigger the acquisition of molecular and phenotypic features that contribute to tumor heterogeneity and therapy resistance (reviewed in (*18*)). Moreover, while stem cells have been denoted as relevant candidates of transformation in certain contexts, progenitors or differentiated cells can also act as tumor-initiating cells by gaining plasticity to change identity (transdifferentiation) and/or re-acquire stem cell-like traits (dedifferentiation) (*18–21*). Among other factors, oncogenic K-Ras (K-Ras^G12D^) was found critical to trigger such early changes of cellular plasticity during MT. *In vitro,* MEF transformation triggered by K-Ras^G12D^, c-Myc and p53 depletion is indeed accompanied by early changes of cellular identity (*14, 22, 23*). *In vivo*, K-Ras^G12D^ alters the identity of alveolar AT2 cells to increase their plasticity and foster the early steps of lung tumorigenesis (*24, 25*) and converts acinar cells into ductal cells in the initial phase of pancreatic adenocarcinoma formation (*26*).

In this context, and even if cellular identity loss and cellular plasticity acquisition are crucial for reprogramming and malignant transformation, the molecular mechanisms coordinating those events and their degree of analogy, remain unknown, mainly because of the lack of flexible genetic tools allowing sophisticated comparative analyses. In this study, we developed “repro-transformable” mice models to compare the early steps of PR – mediated by OSKM – and MT – mediated by oncogenic Ras (K-Ras^G12D^ or H-Ras^G12V^), c-Myc and p53 depletion – in MEF. We identified a combination of somatic markers – namely Bcl11b and Thy1 – that are asynchronously but commonly downregulated during PR and MT. Their combined use enabled the capture of early cellular intermediates (CI) prone or refractory to generate pluripotent or malignant derivatives and the definition of the corresponding cellular roadmaps. Furthermore, comprehensive transcriptomic, epigenomic and functional analyses revealed that these early CI gain plasticity prior to lose significantly MEF identity during both PR and MT. In addition, bio-informatic analyses led to identify the molecular features that are specific or shared by reprogramming and transformation, as well as putative regulators of cellular plasticity. Among those, we found that the basic helix-loop-helix (bHLH) class A TF Atoh8 limits iPS cells generation as well as MT and direct neuron conversion. Mechanistically, we unveiled how Atoh8 broadly constrains cellular plasticity by binding to a specific subset of MEF enhancers and by fine-tuning WNT signalling.

## Results

### A genetic “repro-transformable” model to dissect the early steps of pluripotent reprogramming and malignant transformation

To decipher and compare the early molecular and cellular events of PR and MT, we developed “repro-transformable” mice. OSKM was selected as the prototypical cocktail for PR (*1, 27*) and the cooperation between oncogenic K-Ras (K-Ras^G12D^) and c-Myc was chosen because it was found to trigger cellular identity loss and MT in MEF (*14, 22, 23*). Therefore, R26^rtTA^; Col1a1^4F2A^ mice were crossed with LSL-K-Ras^G12D^; R26^cre-ERT2^ mice and MEF derived (Fig. 1A). The treatment of these “repro-transformable” MEF with doxycycline (Dox) led to the emergence of iPS colonies at an efficiency of 0.21 +/-0.1%. As expected, these cells expressed Nanog and Ssea1 (Fig. S1A) and were able to undergo *in vivo* multilineage differentiation in teratoma (Fig. 1B). In contrast, MT was achieved by tamoxifen (TAM) treatment to induce K-Ras^G12D^ expression (by excision of a Lox-Stop-Lox cassette) and by c-Myc exogenous expression (Fig. 1A). Under these conditions, cells acquired malignant features after serial passaging, as evaluated both *in vitro* and *in vivo.* Foci formation assay indicated the clonal loss c of contact inhibition at an efficiency of 0.66 +/-0.3% (Fig. S1B) while soft agar formation assay showed the acquisition of anchorage-independent growth potential (Fig. S1C). Furthermore, the injection of these cells in Nude mice led to the formation of “liposarcoma-like” tumors (Fig. 1C).

**Figure 1.**
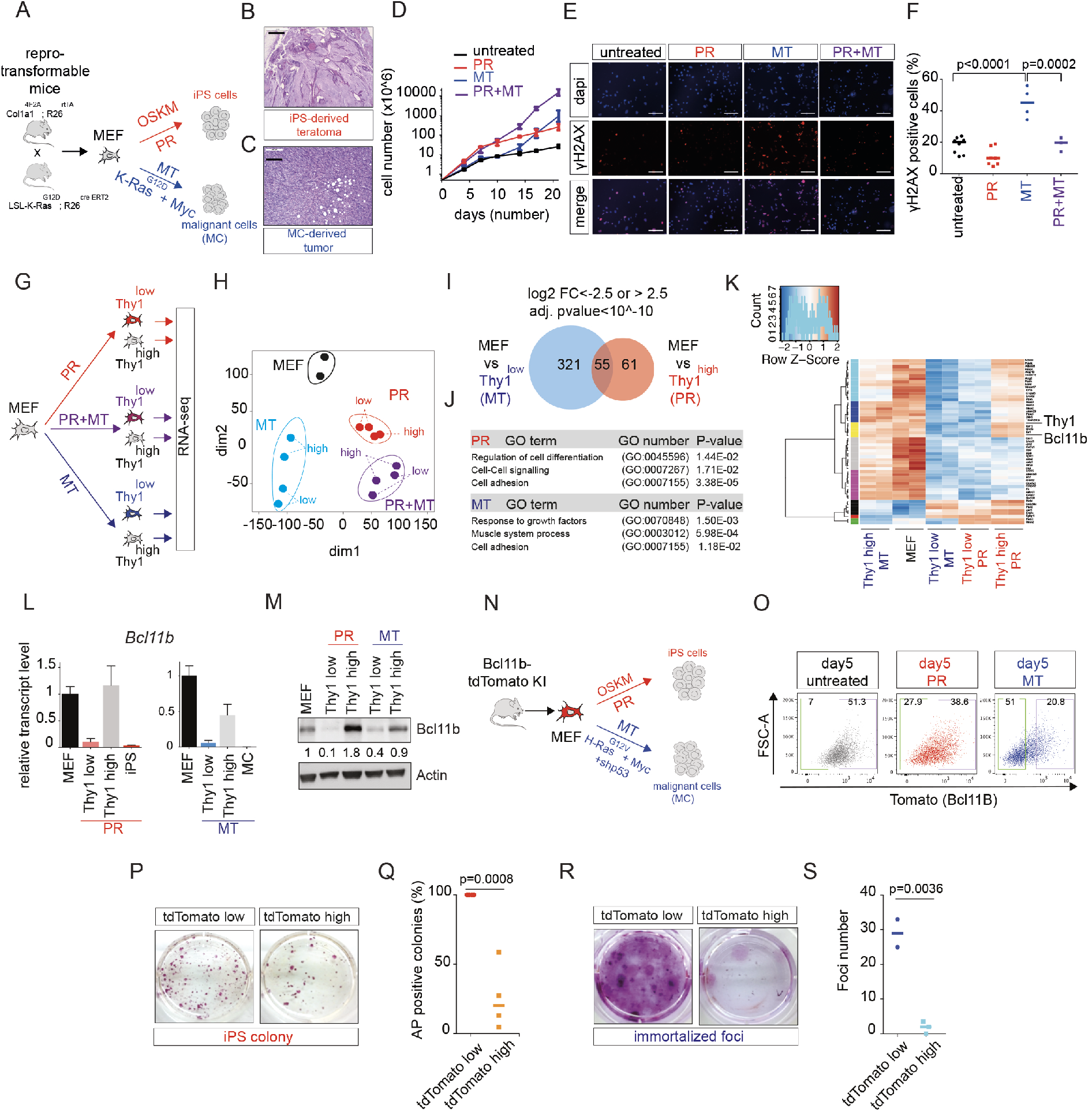
Bcl11b is downregulated in cellular intermediates during both iPS cells generation and malignant transformation. (A) A schematic illustration of the genetic construct of R26^rtTA^; Col1a1^4F2A^; LSL-K-Ras G12D; R26^cre/ERT2^ mice generated to produce MEF. PR (doxycycline-induced OSKM expression) or MT (tamoxifen-induced K-ras^G12D^ expression combined with c-Myc overexpression) are induced to give rise to iPS or malignant cells, respectively. (B) Histological analysis of teratomas derived from PR-induced iPS cells. Scale bar: 1 mm. (C) Tumor formation in nude mice injected with MT-induced malignant cells (MC). Scale bar: 0.2 mm. (D) Proliferation curves of MEF upon PR, MT and PR+MT treatment for 21 days compared to control MEF. n=2 independent experiments. (E) Immunofluorescent staining of PR-, MT-and PR+MT-induced cells for γH2AX after 3 days compared to control MEF. One representative experiment (from three independent experiments). Scale bar: 100 μm. (F) Counting of γH2AX-positive cells depicted in (E). n=3 independent experiments. One-way ANOVA followed by a Tukey’s post hoc test was used. (G) RNA-seq analyses were performed in Thy1^low^ and Thy1^high^ FACS sorted cells after 5 days of PR, MT or the combination of both (PR+MT) compared to control MEF. (H) Principal component analysis. (I) Venn diagram showing the number of genes specifically and commonly regulated in Thy1 ^low^ cells upon PR and MT. (J) Gene Ontology (GO) enrichment for genes differentially expressed in Thy1^low^ cells from PR and MT. (K) Heatmap depicting the expression of the 55 genes commonly regulated in Thy1^low^ cells upon PR and MT. (L) Relative transcript level of *Bcl11b* determined by RT-qPCR in MEF, Thy1 ^low^ and Thy1 ^high^ cells upon PR and MT after 5 days and in iPS cells and malignant cells (MC). n=3 independent experiments. (M) Western blot showing Bcl11b expression in MEF, Thy1^low^ and Thy1^high^ cells upon PR and MT after 5 days. One representative experiment (from three independent experiments). (N) A schematic illustration of the genetic construct of Bcl11b-tdTomato mice to produce MEF. PR (retroviral OSKM expression) or MT (H-ras^G12V^, c-Myc and shp53 viral inductions) are induced to give rise to iPS or MC, respectively. (O) FACS analysis of Bc11b-tdTomato upon 5 days of PR or MT compared to control MEF. (P) Alkaline Phosphatase (AP) staining of iPS colonies generated from tdTomato^low^ and tdTomato^high^ cells FACS-sorted at day 5 of PR. One representative experiment (from four independent experiments). (Q) Counting of AP-positive colonies depicted in (P). n=4 independent experiments. Student t test was used. (R) Foci staining of malignant cells generated from tdTomato^low^ and tdTomato^high^ cells FACS-sorted at day 5 of MT. One representative experiment (from three independent experiments). (S) Counting of foci depicted in (R). n=3 independent experiments. Student t test was used.

The fact that PR and MT can be induced in the same population of “repro-transformable” MEF allowed us to conduct comparative analyses. MEF proliferation was increased by both PR and MT induction but with different kinetics, and this effect was found to be cumulative (Fig. 1D). In contrast, cell cycle features appeared to be specifically modified by MT induction (Fig. S1D). Considering the known action of oncogenes on genome stability, we next evaluated DNA damage after 5 days of reprogramming or transformation. MT induction triggered the formation of H2AX phosphorylation (gH2AX) foci in 45.1+/-10.0% of the cells (Fig. 1E-F). Of note, similar results were obtained with alternative oncogenic triggers including p53 depletion and/or Cyclin E (CCNE) and H-Ras^G12V^ ectopic expression (Fig. S1E-F). In contrast, PR induction did not significantly increase the number of gH2AX foci. Moreover, when PR and MT were simultaneously induced, we found that OSKM significantly prevented gH2AX foci formation triggered by K-Ras^G12D^ and c-Myc (Fig. 1E-F). A preventive effect of OSKM was also observed on apoptosis, as revealed by AnnexinV-PI staining (Fig. S1G-H). We next compared the transcriptomic response to PR, MT and their combination (PR+MT) for 5 days. While PR and MT triggered different transcriptomic changes, their co-induction tends to cluster separately, suggesting a cumulative effect (see principal component analysis in Fig. S1I). Altogether, these results highlight divergent responses of MEF to reprogramming and transformation and a preventive action of OSKM on DNA damage and apoptosis induced by c-Myc and oncogenic K-Ras.

### Identification of somatic markers commonly downregulated in the early steps of pluripotent reprogramming and malignant transformation

With the “repro-transformable” MEF, we next attempted to comparatively track the early cellular changes during PR and MT. Because these early steps are highly inefficient with a small percentage of cells engaging into those paths (*1, 14*), the design of a strategy to capture rare cellular intermediates (CI) was required. In the PR literature, strategies to track CI mainly combined the downregulation of a somatic gene and the activation of a pluripotent one (*3, 28–31*). However, due to the fact that we aimed at tracking early CI of both PR and MT, the use of pluripotent markers was excluded. We therefore attempted to identify a combination of somatic markers downregulated during PR and MT. To do so, we FACS profiled the CD73, CD49d and Thy1 cell surface markers described in the literature (*14, 28, 29, 32*) to assess whether they behave similarly during PR and MT (Fig. S2A). Among those, Thy1 was the sole to be commonly downregulated in a subset of cells after 5 days of both processes (Fig. S2B). We next assessed whether Thy1 downregulation correlated with enhanced reprogramming and transforming potential. For PR, Thy1^low^ and Thy1^high^ cells were FACS sorted after 5 days of OSKM induction and replated at similar densities in reprogramming conditions. Thy1^low^ cells formed significantly more alkaline phosphatase (AP) positive iPS colonies than Thy1 ^high^ cells (2 fold), as previous reported (*3, 28*) (Fig. S2C-D). For MT, a similar FACS sorting strategy (5 days post K-Ras^G12D^/c-Myc induction) was conducted to subject Thy1^low^ and Thy1^high^ cells to foci formation assay. Thy1^low^ cells formed 4-fold more foci than Thy1^high^ cells, indicating that Thy1^low^ cells are more prone to lose contact inhibition (Fig. S2E-F). However, the reprogramming and immortalization efficiencies of Thy1^low^ cells are still very low.

To identify additional markers and refine the strategy, we conducted RNA-seq analysis on untreated “repro-transformable” MEF, Thy1^low^ cells (prone) and Thy1^high^ cells (refractory) sorted by FACS after 5 days of PR, MT and PR+MT (Fig. 1G). PCA showed a sample distribution similar to Fig. S1I, based on PR (dim1) and MT (dim2) induction (Fig. 1H). For each scenario (PR or MT), we next compared untreated MEF and Thy1 ^low^ cells. We identified respectively 116 and 376 genes modulated in Thy1^low^ cells during PR and MT (log2 FC > 2.5 or < −2.5; adjusted p-value < 10^-10^) (Fig. 1I). Statistical over-representation analyses with Pantherdb revealed strong association of the PR-associated genes with “regulation of cell differentiation”, “cell adhesion” and “cell-cell signalling” (Fig. 1J). For MT-associated genes, we found enrichments for genes related to “muscle system process” and mesoderm identity but also “cell adhesion”, similarly to PR. By overlapping both sets of genes, we identified 55 genes commonly regulated in PR-and MT-Thy1 ^low^ cells (Fig. 1K). Among them, we noticed the presence of several genes implicated in cellular adhesion (*Col7a1, Ncam1, Ctgf, Postn*), cancer progression (*Podxl*) and embryonic morphogenesis (*Grem2*). We selected the zinc finger TF Bcl11b for further investigation because it was described as a cellular identity gatekeeper in T cells (*33*). We showed first that Bcl11b expression is high in MEF, specifically decreased in Thy1 ^low^ cells during PR and MT and silenced in iPS and malignant cells (Fig. 1L-M). Of interest, during PR and MT, Bcl11b expression was maintained or even induced in refractory Thy1 ^high^ cells. Next, to assess whether Bcl11b downregulation is correlated with enhanced potential of a cell to become pluripotent or malignant, MEF were derived from Bcl11b-tdTomato reporter knock-in mice (*33*) (Fig. 1N and S2G). FACS analysis confirmed that the majority (90%) of MEF expressed Bcl11b-tdTomato (Fig. 1O). However, after 5 days of PR or MT, we observed the emergence of a subset of Bcl11b^low^ cells (Fig. 1O). For PR, Bcl11b^low^ cells, sorted by FACS after 5 days of OSKM induction, formed on average 7-fold more AP+ iPS colonies than their Bcl11b^high^ counterparts (Fig. 1P-Q). For MT, Bcl11b^low^ cells formed immortalized foci at a 10-fold higher efficiency than Bcl11b^high^ (Fig. 1R-S). Altogether, these results identified Bcl11b as a faithful indicator of the ability of cells to engage into pluripotency and immortalization paths.

### The combined downregulation of Bcl11b and Thy1 delineates an early plastic cellular intermediate, highly amenable to reprogramming and transformation

The previous findings prompted us to investigate whether the combined downregulation of Bcl11b and Thy1 allows to capture early plastic CI. First, interrogation of single cell RNA-seq (sc-RNA-seq) data conducted during MEF reprogramming confirmed *Bcl11b* and *Thy1* downregulation but also revealed different behaviors of the 2 transcripts. While *Thy1* was homogeneously repressed rapidly after 2 days of PR, as previously described (*3*), *Bcl11b* was still heterogeneously expressed at day3, indicating that both factors are not regulated similarly and delineate different subpopulations (Fig. S3A) (*11*). Therefore, we profiled Bcl11b and Thy1 changes during PR (induced by OSKM) and MT (induced by H-Ras^G12V^, c-Myc and p53 depletion) by FACS (Fig. 2A). To begin the experiments with an homogeneous cell population, Bcl11b-tdTomato^high^/Thy1^high^ MEF (representing 90% of the population) were FACS sorted to purity. In the absence of reprogramming or transformation, sorted MEF stably maintained a Bcl11b-tdtomato^high^/Thy1^high^ phenotype in culture (Fig. S3B). By day 17, most of the cells displayed downregulation of both proteins, similarly to iPS and malignant cells, as expected (Fig. 2A). Interestingly, we next demonstrated that the rare Bcl11b^low^/Thy1 ^low^ cells, that emerged by day3 of PR and MT, harbored a strong potential to reprogram or transform, when compared to their Bcl11b^high^/Thy1 ^high^ respective counterparts. For PR, Bcl11b^low^/Thy1 ^low^ cells (hereafter entitled PRP for Pluripotent Reprogramming Prone) formed 13-fold higher iPS colonies than Bcl11b^high^/Thy1 ^high^ cells (hereafter entitled PRR for Pluripotent Reprogramming Refractory) (Fig. 2B-C). For MT, Bcl11b^low^/Thy1^low^ cells (hereafter entitled MTP for Malignant transformation Prone) formed foci at a 4-fold higher rate than Bcl11b^high^/Thy1 ^high^ cells (hereafter entitled MTR for Malignant transformation Refractory) (Fig. 2D-E). These data indicate that the combined use of Thy1 and Bcl11b allowed the isolation of early CI highly amenable to form pluripotent or immortalized derivatives.

**Figure 2.**
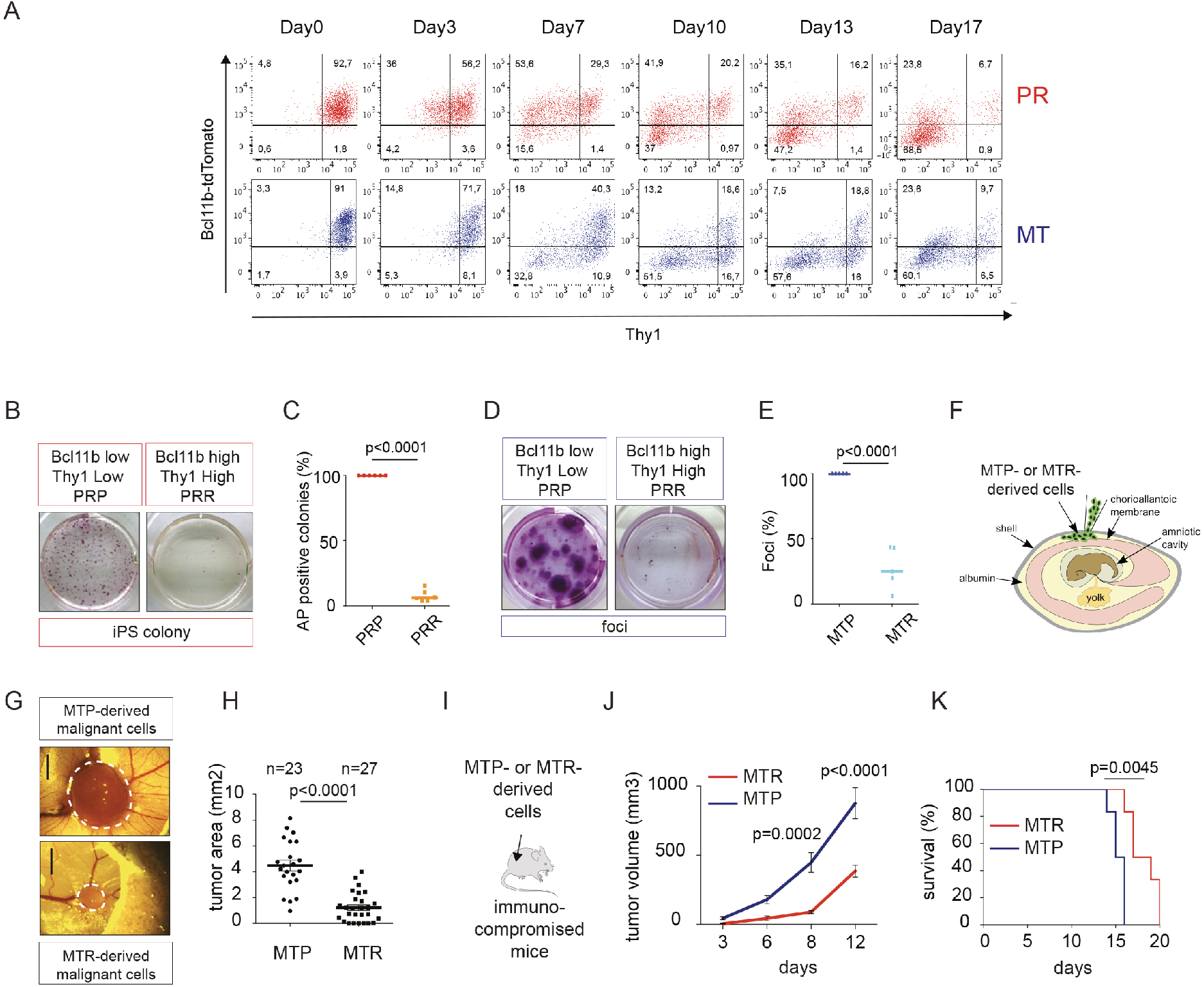
The combined downregulation of Bcl11b and Thy1 delineates a cellular intermediate highly amenable to reprogramming and transformation. (A) FACS profile of Bcl11b-tdTomato and Thy1 upon PR and MT from day 0 to day 17. (B) Alkaline Phosphatase (AP) staining of iPS colonies generated from Bcl11b-tdTomato^Low^/Thy1^Low^ (PRP: Pluripotent Reprogramming Prone) and Bcl11b-tdTomato^High^/Thy1^High^ (PRR: Pluripotent Reprogramming Refractory) cells. The subpopulations were FACS-sorted at day 7 of PR, replated at equal densities in PR conditions and AP staining was performed after 10 days. One representative experiment (from six independent experiments). (C) Counting of AP-positive colonies depicted in (B). n=6 independent experiments. Student t test was used. (D) Foci staining of malignant cells generated from Bcl11b-tdTomato^Low^/Thy1^Low^ (MTP: Malignant transformation Prone) and Bcl11b-tdTomato^High^/Thy1^High^ (MTR: Malignant transformation Refractory) cells. The subpopulations were FACS-sorted at day 7 of MT, replated at equal densities in MT conditions and foci staining was performed after 15 days. One representative experiment (from five independent experiments). (E) Counting of foci depicted in (D). n=5 independent experiments. Student t test was used. (F) Scheme of the chicken chorioallantoic membrane (CAM) assay. MTP- or MTR-derived cells were inoculated on the CAM in the egg of chick embryos at E11. (G) Pictures of tumors generated by MTP- and MTR-derived cells. The size of the tumor was evaluated after 7 days. White dotted lines delineate the size of the tumors. (H) Counting of the tumors of (G). Student’s t-test was used, and two-sided p-values are indicated. (I) Scheme of the xenograft assay. MTP- or MTR-derived cells were injected subcutaneously into immunocompromised SCID mice. (J) Tumor growth curves. Tumor size was monitored every 3 days until day 12. 2-way ANOVA was used (K) Survival curve of mice injected with MTP or MTR-derived malignant cells. Log-rank test was used.

We next assessed whether the early downregulation of Bcl11b and Thy1 had consequences on the subsequent acquisition of pluripotent and malignant features. For PR, iPS cell lines, established from PRP or PRR cells sorted after 5 days of reprogramming, expressed similar levels of Oct4, Nanog and Sox2. This result indicates that the kinetic of Thy1 and Bcl11b downregulation does not broadly impact the establishment of pluripotent features, in agreement with reports describing that iPS cells emerging from different routes and cells-of-origin are strikingly similar (data not shown)(*9*). However, because the status of the cell-of-origin can have a profound impact on the molecular and phenotypic features of a malignant outcome (*34, 35*), we conducted similar approaches during MT. To do so, we FACS sorted MTP and MTR cells after 5 days of MT, replated them at equal densities and established independent polyclonal cell lines by serial passaging. While MTP- and MTR-derived cell lines presented similar growth curves *in vitro* in 2D (Fig. S3C), we found that 3 independent MTP-derived lines formed colonies in soft agar at a 7-fold higher efficiency than MTR-derived ones, indicating a higher potential for anchorage-independent growth (Fig. S3D-E). We next employed the chick chorioallantoic membrane (CAM) assay as an *in vivo* model of tumor development (Fig. 2F). GFP-labelled MTR- and MTP-derived cell lines were seeded on the upper CAM of day11 chick embryos and the surface area of the primary tumors calculated 7 days later. The size of the GFP+ tumors generated with MTP-derived cells was significantly higher than those generated with MTR-derived cells (Fig. 2G-H and S3F), indicating an accelerated tumor growth *in vivo*. To reinforce this finding, we performed xenograft assays by injecting MTR- and MTP-derived cells into immunocompromised mice (Fig 2I). The growth of MTP-derived tumors was significantly faster than MTR-derived ones (Fig. 2J) and the mice survival rate significantly reduced (Fig. 2K). Altogether these data indicate that the early and combined loss of Bcl11b and Thy1 delineates the emergence of plastic CI, highly amenable to form iPS cells during PR and highly tumorigenic cells during MT.

### The sequential downregulation of Bcl11b and Thy1 allows to define the comparative roadmaps of reprogramming and transformation

Using Bcl11b and Thy1, we next sought to define the cellular roadmaps of reprogramming and transformation. To demonstrate that the Bcl11b/Thy1 profile changes observed in Fig. 2A reflected the transition of individual cells from one stage to the next, and not merely the loss of one major population and expansion of another minor population, each fraction was sorted, replated for 48 hours in culture before being re-analysed by FACS. For PR, the 4 following subpopulations were FACS sorted after 7 days of reprogramming: Bcl11b^high^/Thy1^high^ (PRR), Bcl11b^low^/Thy1^high^ (PR1 = Pluripotent Reprogramming Intermediate 1), Bcl11b^high^/Thy1^low^ (PR2 = Pluripotent Reprogramming Intermediate 2), Bcl11b^low^/Thy1^low^ (PRP) (Fig. 3A). The progression of cellular fates and the corresponding transition rates revealed the diverse routes triggered by OSKM. First, we observed that the PRP state, characterized by the combined downregulation of Bcl11b and Thy1, is stable. Indeed, FACS sorted PRP cells did not transit efficiently into other states (Fig. 3A – top left panel). Secondly, we found that the PRR and PR2 cells generated PRP cells at very low rate while PR1 cells transit into PRP very efficiently (35%). This suggests that reaching the PR1 state, characterized by Bcl11b downregulation, is a rate-limiting step of iPS cell generation. Importantly, these cellular progressions were correlated with the relative capacities of the corresponding populations to form AP-positive colonies (Fig. 3B-C). When a similar analysis was conducted with MT, we observed that the MTP state is also quite stable (Fig. 3D – top left panel). However, the MTR and MT1 cells were poorly efficient at generating MTP while MT2 cells did transit toward this state at high efficiency, indicating that Thy1 downregulation could constitute a rate-limiting step, in agreement with the relative ability of these subpopulations to form immortalized foci (Fig. 3E-F). On the basis of total cell numbers in each gate and their potential to reprogram/immortalize, we generated the cellular roadmaps presented in Figure 3G. Owing to the Bcl11b/Thy1-based strategy, we revealed hierarchical steps and cellular transition states during PR and MT.

**Figure 3.**
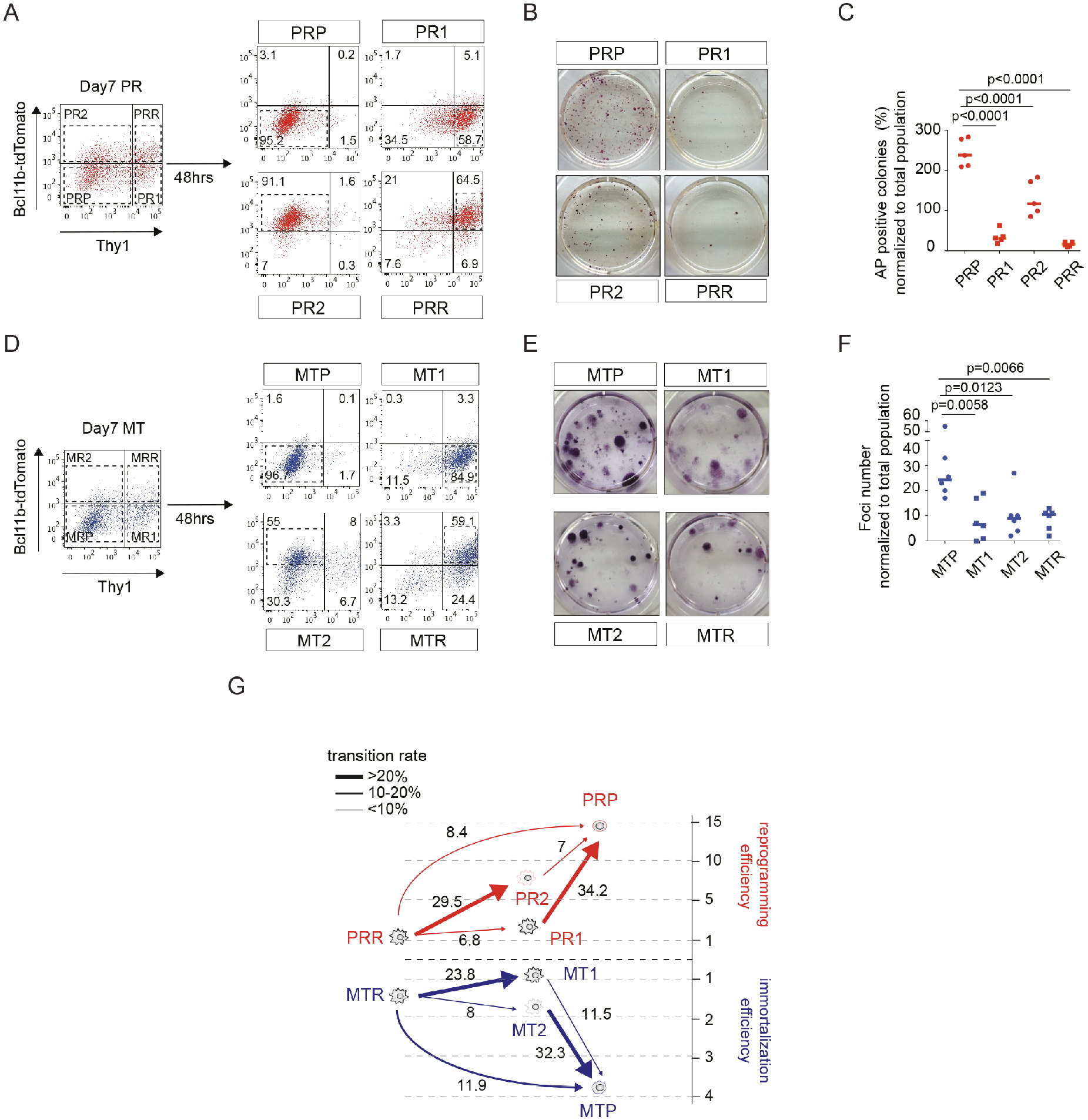
The sequential downregulation of Bcl11b and thy1 delineates cellular roadmaps toward pluripotency and malignancy. (A) Four different subpopulations (Bcl11b-tdT omato^high^/Thy1^high^, Bcl11b-tdT omato^high^/Thy1^low^, Bcl 11b-tdT omato^low^/Thy1^high^ and Bcl11b-tdTomato^low^/Thv1^low^) were FACS-sorted at day 7 of PR and replated. After 2 days, the expression profile of Bcl11b and thy1 were analyzed for the 4 subpopulations by FACS. (B) AP staining of iPS colonies generated from the different subpopulations described in (A) FACS-sorted at day 7 of PR. One representative experiment (from five independent experiments). (C) Counting of AP-positive colonies depicted in (B). n=5 independent experiments. One-way ANOVA followed by a Tukey’s post hoc test was used. (D) 4 different subpopulations (same as in (A)) were FACS-sorted at day 7 of MT and replated. After 2 days, the expression profile of Bcl11b and Thy1 were analyzed for the 4 subpopulations by FACS. (E) Foci staining of malignant cells generated from the different subpopulations described in (D) FACS-sorted at day 7 of MT. One representative experiment (from six independent experiments). (F) Counting of foci depicted in (E). n=5 independent experiments. One-way ANOVA followed by a Tukey’s post hoc test was used. (G) A schematic illustration of the trajectories taken by the different subpopulations of cells during PR and MT and the reprogramming/transforming efficiencies associated. PRP: pluripotent reprogramming prone, PR1: pluripotent reprogramming intermediate 1, PR2: pluripotent reprogramming intermediate 2, PRR: pluripotent reprogramming refractory, MTP: malignant transformation prone, MT1: malignant transformation intermediate 1, MT2: malignant transformation intermediate 2, MTR: malignant transformation refractory.

### Comprehensive analysis of chromatin accessibility reconfigurations in CI during pluripotent reprogramming and malignant transformation

We next investigated the reconfigurations of chromatin accessibility in the CI defined in Figure 3G by conducting ATAC-seq on FACS-sorted cells after 5 days of PR and MT. Untreated MEF, iPS and malignant cells generated respectively by PR and MT were included (Fig. 1A-C). PCA analysis showed that PRP and MTP cells, respectively prone to generate iPS and malignant cells, segregated together on the X-axis (dim1) (Fig. 4A) and towards the direction of the fully reprogrammed/transformed cells (Fig. S4A), suggesting the existence of common chromatin accessibility changes in these CI. To test this, we classified the chromatin peaks in clusters defining regions that (i) were accessible in MEF but exhibit loss of accessibility in both PRP/MTP over MEF (Cluster 1, n=4522); (ii) became accessible in both PRP/MTP over MEF (Cluster 2, n=3015); (iii) were specifically lost (Cluster 3, n=3776) or gained (Cluster 4, n=5464) in PRP over MEF and MTP or (iv) were specifically lost (Cluster 5, n=13245) or gained (Cluster 6, n=17282) in MTP over MEF and PRP (Fig. 4B). This clustering highlighted that the number of peak modifications shared by PRP and MTP (C1+C2=7537) is closed to the PRP-specific modifications (C3+C4=9240) but lower than the MT-specific changes (C5+C6=30527). These results also indicated that the early steps of PR and MT trigger similar and specific changes in chromatin accessibility. Next, analysis of DNA motif enrichment in the ATAC-seq clusters revealed different families of TF (Fig. 4C and S4B). Among those, we found that a subset of regulatory elements losing accessibility specifically in PRP and MTP were enriched for the FosL1 motif (C3 and C5) meanwhile another subset of FosL1-containing peaks were gained commonly in PRP and MTP (C2), suggesting shared and specific relocation of this factor during PR and MT. We next assessed whether Fosl1 functionally regulates both processes. FosL1 depletion (Fig. S4C, 52.7% average knockdown efficiency) led respectively to a 4-fold reduction in the number of immortalized foci in MT (Fig. 4D-E) and an averaged 6-fold increase in reprogramming efficiency during PR (Fig. 4F-G), highlighting antagonistic functions. Altogether, these epigenomic analyses depict common and specific chromatin accessibility changes during PR and MT and unveil FosL1 as a functional regulator.

**Figure 4.**
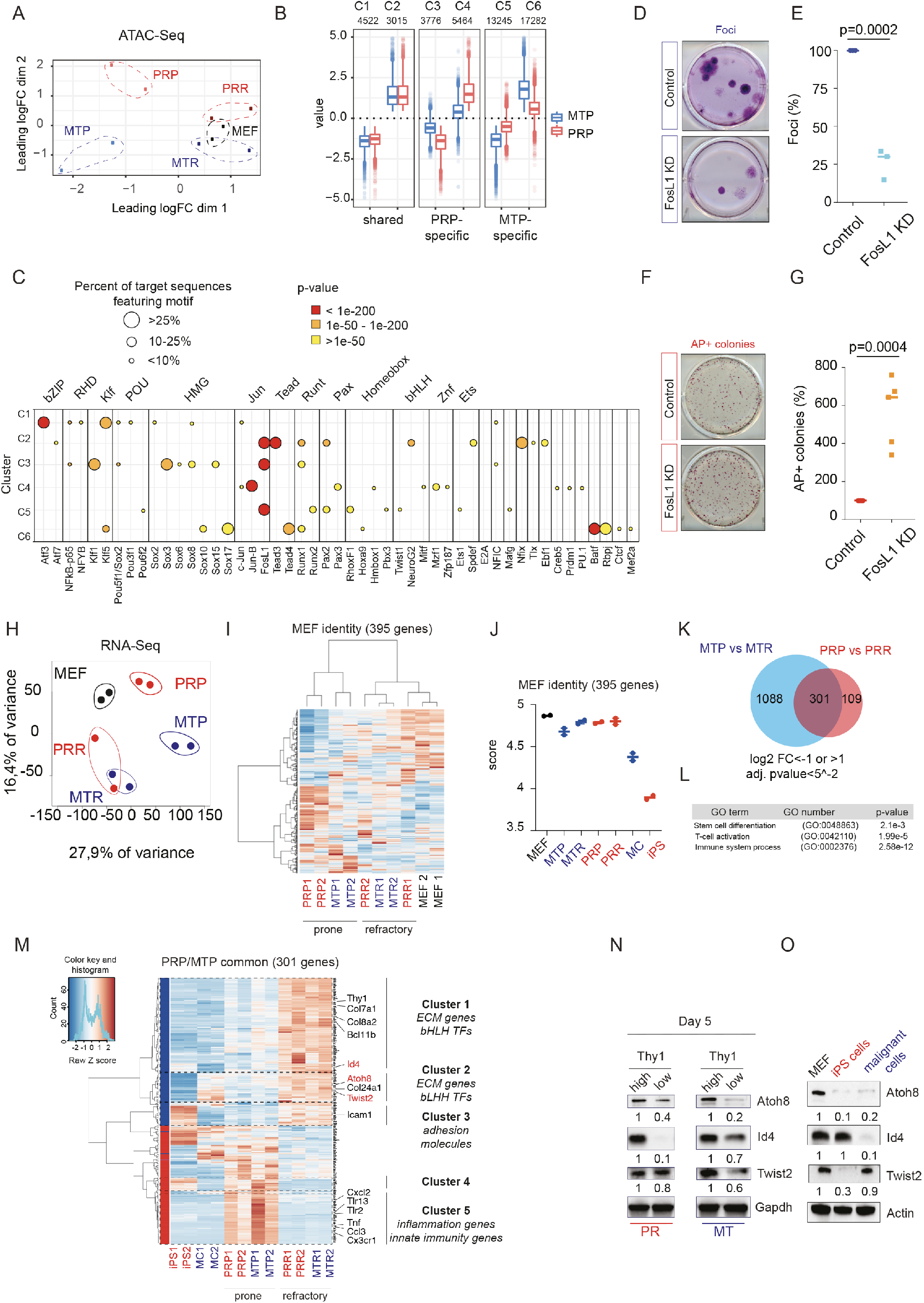
Epigenomic and transcriptomic reconfiguration in cellular intermediates during reprogramming and transformation. (A) Principal component analysis conducted on ATAC-seq analysis of MEF, PRP, PRR, MTP and MTR samples depicted in Fig. 3G. (B) Definition of clusters in ATAC-seq regions (see main text for description). (C) Enrichment in transcription factors motifs in ATAC-seq clusters. Each point represents a significant enrichment in the motif (x axis) for the cluster (y axis). Point size represents the proportion of sequences in the cluster featuring the motif and color gradient the enrichment significance. (D) Foci staining of immortalized cells generated in control- or FosL1-depleted settings. One representative experiment (from three independent experiments). (E) Counting of foci depicted in (D). n=3 independent experiments. Student t test was used. (F) Alkaline Phosphatase (AP) staining of iPS colonies generated control- or FosL1-depleted settings. One representative experiment (from six independent experiments). (G) Counting of AP-positive colonies depicted in (F). n=6 independent experiments. Student t test was used. (H) Heatmap clustering the different samples based on the expression of the 395 genes constituting the MEF identity score defined in (*9*) using RNA-seq data from MEF, PRP, PRR, MTP and MTR samples depicted in Fig. 3G. (I) Ssgsea analysis of the MEF identity score in similar samples as (H). The iPS and malignant cells (MC) datasets were included (see Methods for details). (J) Principal component analysis of normalized gene expression of similar samples as (H). (K) Venn diagram showing the number of genes specifically and commonly regulated in prone cells (PRP and MTP) upon PR and MT compared to the refractory cells (PRR and MTR). (L) Statistical overrepresentation assays conducted with Pantherdb on genes differentially expressed in PRP and MTP cells. (M) Heatmap clustering the genes based on the selection of 301 commonly deregulated genes from (K). (N) Western blot of Atoh8, Id4 and Twist2 in Thy1^low^ and Thy^high^ subpopulations at day 5 of PR or MT. (O) Western blot of Atoh8, Id4 and Twist2 in MEF, iPS and malignant cells.

### Comprehensive analysis of transcriptome reconfigurations in CI during pluripotent reprogramming and malignant transformation

We next conducted RNA-seq on the same samples as ATAC-seq. PCA revealed first, and similarly as ATAC-seq, that PRP and MTP segregated together on the X-axis, suggesting common transcriptomic changes (Fig. 4H). We next exploited published datasets to characterize the PRP and MTP CI. We first focused on cellular identity using a MEF signature of 395 genes previously defined by the Mogrify algorithm (*9, 36*). Even if the PRP and MTP CI clustered together in a heatmap analysis (Fig. 4I), their identity score was nearly similar to MEF and refractory PRR/MTR, and therefore significantly higher than iPS and malignant cells (Fig. 4J). This finding is consistent with the fact that PRP and MTP isolation is exclusively based on the downregulation of somatic markers, in contrast to previously published intermediates that were sorted based on the reactivation of the pluripotent marker Ssea1 (*5*). Similarly, PRP and MTP cells did not present elevated levels of *CD73* and *CD49d,* that were found to delineate another class of reprogramming intermediates (Fig. S4D) (*30*). Finally, based on a recent report from Lander and colleagues, we showed that PRP and MTP cells harbored significant reductions in some stromal markers, such as *Csf1, Prrx1* and *Id3,* confirming that that they just begun to repress this identity. However, most of the early MET markers (*Fut9, Zic3, Dmrtc2* and *Pou3f1*) were not induced (except *Shisa8),* reinforcing the view that the PRP and MTP CI have not yet engaged into a MET trajectory (Fig. S4E)(*10*). Altogether, these analyses indicate that the gain of cellular plasticity of the PRP and MTP CI is not correlated with a significant loss of cellular identity or an engagement into a particular MET state.

Because of this finding, we next sought to define the transcriptomic features of the PRP and MTP CI. Volcano plot showed that 410 genes were differentially expressed between PRP (prone) and PRR (refractory) and 1389 genes between MTP (prone) and MTR (refractory), with a common signature of 301 genes commonly deregulated in PRP and MTP (170 down and 131 up, adjusted p-value < 5.10^-2^; log2 FC >1 or <-1) (Fig. 4K). Statistical overrepresentation assays using Pantherdb revealed enrichments in stem cell differentiation but also unexpectedly immunity (Fig. 4L). Next, we conducted a heatmap analysis including iPS and malignant cells datasets to define clusters of genes permanently or transiently regulated in CI during PR and/or MT (Fig. 4M). Cluster 1 corresponds to genes permanently repressed during PR and MT (downregulated in PRP, MTP, iPS and malignant cells when compared with PRR and MTR), including *Thy1* and *Bcl11b,* as expected, but also *Col24a1, Col4a6* and *Col7a1,* possibly reflecting a reorganization of the extracellular matrix (*37*). Clusters 2 and 3, that encompass genes transiently repressed in prone CI but reactivated respectively in malignant or iPS cells, highlighted cell adhesion molecules such as *Icam1,* previously identified in PR intermediates (*29*). We also identified a restricted cluster 4 of 14 genes transiently induced in prone CI but re-expressed in iPS cells. Interestingly, cluster 5, that regroups genes transiently induced in PRP and MTP CI, was highly enriched in genes associated with innate immunity and inflammation (*Tnf, Ccl3, C3ar1, C5ar1, Cxcl2, Tlr13, Tlr2*) In addition, a pantherdb protein class analysis revealed a striking downregulation of a network of basic loop-helix-loop (bHLH) TFs, encompassing *Atoh8, Id4* and *Twist2,* in clusters 1-2 (Fig. 4M). Because BHLH family members were described to play important roles in development, oncogenesis (*38, 39*) and in the control of cellular identity (*40–43*), we conducted further investigations. At the protein level, we first confirmed Atoh8, Id4 and Twist2 downregulation in CI after 5 days of PR and MT (Fig. 4N). However, Id4 and Twist2 proteins were re-expressed in iPS and malignant cells, respectively, while Atoh8 silencing was maintained (Fig. 4O). Altogether, these results revealed the existence of common transcriptomic changes in CI during PR and MT, encompassing a transient induction of immune and inflammation-related genes and a permanent downregulation of the bHLH TF Atoh8.

### Atoh8 constrains mouse iPS cells generation by hindering the reactivation of the pluripotent network

Interrogation of published datasets confirmed Atoh8 downregulation in later CI during PR (Fig. S5A), but also showed accumulation of the repressive histone mark H3K27Me3 on its promoter (Fig. S5B) (*3, 5, 9*), in adequation with its lack of expression during mouse pre-implantation development (Fig. S5C)(*44*). Atoh8 RNAi-mediated knockdown (KD), prior to PR induction (Fig. 5A, S5D-E, 77.7% average KD efficiency), led to a 4-fold increase of PR efficiency, as evaluated by alkaline phosphatase-positive (AP+) (Fig. S5F-G) or Pou5f1-GFP-positive (Pou5f1-GFP+) colonies counting (Fig. 5B-C). Similar increases of PR efficiency were observed by targeting Atoh8 locus with two independent CRISPR/Cas9 guides (Fig. S5H-K), demonstrating that Atoh8 constrains mouse iPS cells generation. To assess whether Atoh8 depletion affects the acquisition of pluripotent features, independent control and Atoh8-KD iPS cell lines were established. Atoh8-KD iPS cell lines were found to express similar Oct4, Sox2 and Nanog levels as control cells (Fig. S5L-M) and to differentiate into the three germ layers in teratoma when injected into immunocompromised mice (Fig. 5D). During PR, reprogramming cells turn on the endogenous pluripotency network and become independent from the OSKM transgene (*28*). By conducting kinetic analyses, we noticed that the emergence of Pou5f1-GFP+ cells, detected as early as day4 after OSKM induction, is significantly accelerated by Atoh8 depletion (Fig. 5E-F), indicating that this bHLH TF constrains the pace of the reactivation of the endogenous Oct4 locus. To evaluate the acquisition of transgene independency, OSKM Dox-inducible Pou5f1-GFP reporter MEF were exposed to Dox for a short period (6 days) and iPS cell colonies emergence monitored by day 15 (Fig. 5G). In these stringent settings, Atoh8-KD cells succeeded to form AP+ iPS colonies while control cells failed, demonstrating that Atoh8 depletion accelerates the emergence of transgene-independent cells during PR (Fig. 5H-I). Pou5f1-GFP+ iPS cell lines derived from Atoh8-KD cells sorted at day 6 of PR expressed similar levels of Oct4, Nanog and Sox2 as *bona fide* iPS lines (Fig. 5J and S5N-O). Altogether, these results showed that Atoh8 hinders rodent iPS cells generation by limiting the reactivation of the endogenous Pou5f1 and the acquisition of transgene independency.

**Figure 5.**
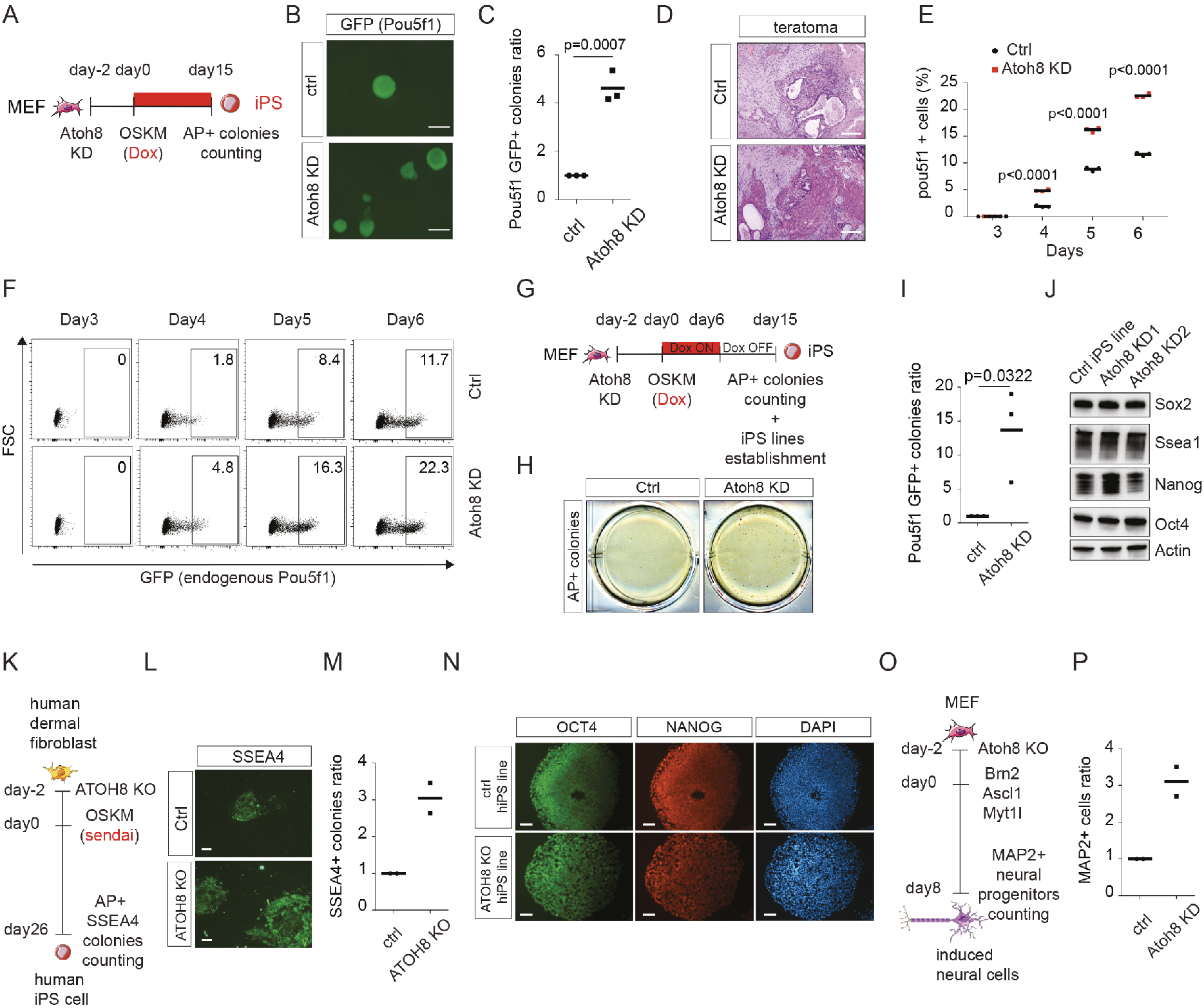
Atoh8 is a broad-range gatekeeper of cellular identity during reprogramming and transdifferentiation. (A) Experimental scheme. Cells were infected with lentiviral shRNA particles targeting control or Atoh8 sequences. 48 hours later, pluripotent reprogramming was induced by dox treatment (OSKM expression). iPS colonies were scored at day 15 by alkaline phosphatase (AP) staining and/or Pou5f1 GFP microscopic observation. (B) Picture representing Pou5f1-GFP+ colonies at day 15 of PR in control and Atoh8 KD conditions, representative of three independent experiments. Scale bar: 100μm (C) Pou5F1+ colony counting. Data are the mean ± s.d. (n=3 independent experiments). Student’s t-test was used, and two-sided p-values are indicated. (D) Pictures depicting histological analysis of teratomas derived from control and Atoh8 KD iPS cell lines. 2 independent teratoma were analyzed per cell line. (E) Graph depicting Pou5f1-GFP+ percentage of cells during reprogramming. Squares correspond to control, circles to Atoh8 KD settings. Data are the mean ± s.d. (n=3 independent experiments). Student’s t-test was used, and two-sided p-values are indicated. (F) FACS analysis showing the emergence of Pou5f1-GFP positive cells from day 3 to day 6 of pluripotent reprogramming performed in control- and Atoh8 KD-settings. (G) Scheme depicting the experimental design. Cells were infected with lentiviral shRNA particles targeting control or Atoh8 sequences. PR was induced by dox treatment for 6 days, then cells were harvested in KSR+Lif medium without dox for the remaining 9 days. iPSC colonies were scored at day 15 by AP+ staining. (H) Picture representing AP+ colonies at day 15 of reprogramming in control and Atoh8 KD settings, representative of three independent experiments. (I) Colony counting. Data are the mean ± s.d. (n=3 independent experiments). Student’s t-test was used, and twosided p-values are indicated. (J) Western blot of Sox2, Ssea1, Nanog and Oct4 in control and 2 independent iPS cell lines obtained from accelerated reprogramming (Atoh8 KD1 and KD2). Scale bar: 100μm. (K) Experimental scheme of human pluripotent reprogramming. Cells were infected with lentiviral sg#control and sg#Atoh8 particles. 48 hours later, pluripotent reprogramming was induced with OSKM Sendai viruses and iPS colonies scored at day 26 by AP+ staining and SSEA4+ imaging. (L) Picture representing SSEA4+ colonies at day 26 of reprogramming in control and Atoh8 KO settings, representative of two independent experiments. (M) Colony counting. Data are the mean ± s.d. (n=2 independent experiments). Scale bar: 150μm. (N) Immunofluorescence for Oct4 and Nanog in same cells as (L). (O) Scheme depicting MEF to neuron transdifferentiation. Cells were infected with lentiviral control and Atoh8 targeting particles. 48 hours later, MEF to neuron reprogramming was induced through lentiviral infection of Brn2, Ascl1 and Mtyl1. Cells were changed from MEF to N3 medium at day 3 and induced neurons iNs scored at day 8 by MAP2 immunofluorescence. (P) iNS counting per field. Data are the mean ± s.d. (n=2 independent experiments).

### Atoh8 constrains cellular plasticity during human pluripotent reprogramming and transdifferentiation

To broaden the role of Atoh8 as a general obstacle to cellular plasticity acquisition, we assessed its function in human dermal fibroblast (HDF) reprogramming during which it was also found to be rapidly downregulated (Fig. S6A) (*45*). CRISPR/Cas9-mediated Atoh8 KO in HDF, prior to induce iPS cells generation with OSKM, significantly improves PR efficiency, as determined by AP+ staining (Fig. S6B-C) or SSEA4 live immunofluorescence (Fig. 5K-M). Control and Atoh8-KO iPS cell lines were found to express comparable levels of pluripotency markers in self-renewing conditions (Fig. 5N and S6D), as well as multilayered differentiation genes during embryoid bodies (EB) formation (Fig. S6E). We next assessed whether Atoh8 limits cellular plasticity during MEF to neuron transdifferentiation induced by the TFs Brn2, Ascl1 and Myt1l (BAM) (Fig. 5O) (*42*). Atoh8 depletion led to a 3-fold increase in the number of MAP2+ induced neuronal cells (iN), as revealed by immunofluorescence (Fig. 5P and S6F). Altogether, these data demonstrated that Atoh8 broadly constrains the acquisition of cellular plasticity during human and mouse TF-mediated cell conversions.

### Atoh8 constrains malignant transformation and cancer cell phenotypic plasticity

We next wondered if Atoh8 functionally interferes with cellular plasticity during immortalization and transformation. First, when MT was induced in MEF by combining c-Myc and K-Ras^G12D^ expression with p53 depletion, we found that Atoh8 depletion significantly increased the ability of cells to form immortalized foci and grow independently of anchorage (Fig. 6A-C and S7A-C). Comparable results were obtained using two independent CRISPR/Cas9 guides (S7D-G), demonstrating that Atoh8 constrains the efficiency of immortalization and transformation. Next, to assess whether the pace at which cells acquire malignant properties is controlled by Atoh8, Control and Atoh8-KD MEF were induced for MT and subjected to soft agar assays as early as 6 days later (Fig. 6D). In these stringent settings, Atoh8-KD cells formed colonies while control cells barely failed (Fig. 6E-F), demonstrating that Atoh8 constrains the acquisition of anchorage-independent growth properties. In line with a function for Atoh8 in preventing tumorigenesis, TCGA analyses revealed a significant downregulation of *Atoh8* expression in malignant tissues when compared with paired peritumoral ones in various cancer types including lung adenocarcinoma and squamous cell carcinoma together with breast cancer and prostate adenocarcinoma (Fig. 6G)(*46*).

**Figure 6.**
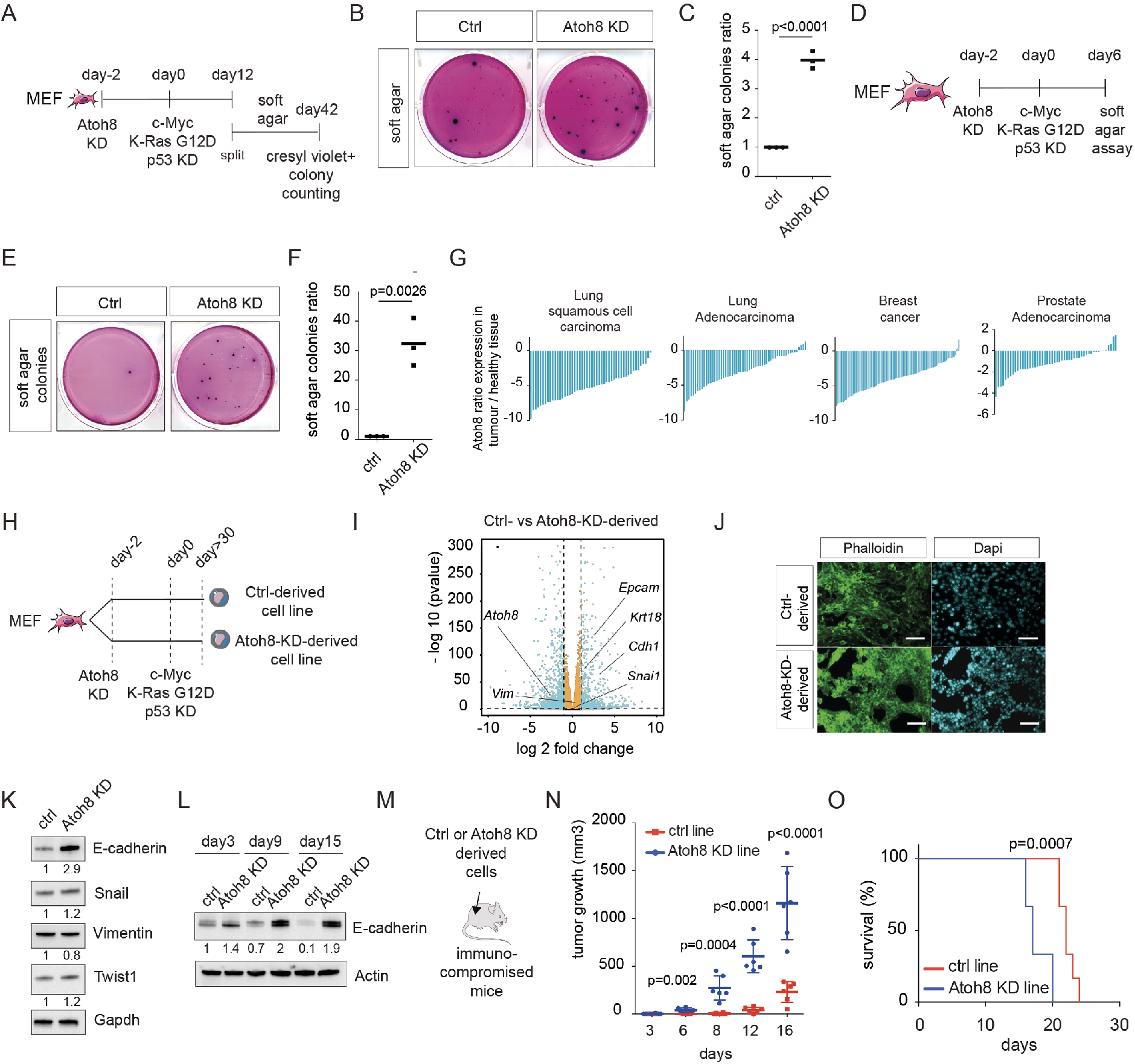
Atoh8 constrains malignant transformation and cancer cell phenotypic plasticity. (A) Scheme depicting transformation assays. Cells were infected with lentiviral particles targeting control and Atoh8 sequences. 48 hours later, malignant transformation was induced by combining 4-OHT treatment (to induce K-RasG12D expression), lentiviral shRNA particles targeting p53 and retroviral particles inducing c-Myc exogenous expression. Cells were split three times and soft-agar colonies were scored after 30 days by Cresyl-violet staining. (B) Picture representing soft-agar colonies in control and Atoh8 KD settings, representative of three independent experiments. (C) Colony counting. Data are the mean ± s.d. (n=3 independent experiments). Student’s t-test was used, and two-sided p-values are indicated. (D) Experimental scheme used to evaluate Atoh8 function in the pace of transformation. Cells were infected with lentiviral particles targeting control and Atoh8 sequences. 48 hours later, MEF transformation was induced as described in (A). Soft agar assays were conducted after 6 days and colonies scored at day 30 by Cresyl-violet staining. (E) Picture representing colonies in control and Atoh8 KD settings, representative of three independent experiments. (F) Colony counting. Data are the mean ± s.d. (n=3 independent experiments). Student’s t-test was used, and two-sided p-values are indicated. (G) Histogram depicting *Atoh8* transcript levels in patients. Data are presented as a log2 of the ratio of Atoh8 FPKMs between malignant and healthy tissues in paired samples. (H) Scheme depicting polyclonal the establishment of control- and Atoh8-KD-derived cell lines. Cells were split for at least 10 passages (around 30 days) before subsequent analyses. (I) Volcano plot comparing transcriptome of control- and Atoh8-KD-derived cell lines. Each dot corresponds to a transcript. Blue dots present a log2 fold change major than 1 or minor than −1, and adjusted p-value inferior to 0.00001. (J) Phalloidin immunofluorescence of control- and Atoh8-KD-derived cell lines. Scale bar: 80μm. (K) Western blot of Cdh1, Snail, Vim and Twist1 cell lines from (H). (L) Western blot of Cdh1 in control and Atoh8 KD settings of malignant transformation at passage 1, 3 and 5 (day3, day9 and day15). (M) Scheme depicting mouse xenograft assays. (N) Xenograft tumor volume over time after injection of cell lines from (H). Data are the mean ± s.d. (n=6 independent mice per group). Student’s t-test was used, and two-sided p-values are indicated. (O) Survival growth of mice from (M). Data are the mean ± s.d. (n=6 independent mice). Kaplan Meyer test was used.

While the consequences of Atoh8 loss on established cancer cells has been addressed for example in hepatocellular carcinoma, its role in the acquisition of phenotypic plasticity during transformation remains unknown (*35, 47*). To test it, we established transformed cell lines by combining c-Myc and K-Ras^G12D^ expression with p53 depletion in presence (Ctrl) or absence (Atoh8-KD) of Atoh8, as depicted in Figure 6H. RNA-Seq conducted on these Ctrl- and Atoh8-KD-derived cell lines led to identify 803 differentially expressed genes (log2 FC > 1 and < 1; adjusted pvalue < 10^-5^) (Fig. 6I), indicating the profound consequences of Atoh8 depletion on the acquisition of malignancy. Statistical overrepresentation assay using Pantherdb showed an enrichment for genes related to “cell adhesion” (Fig. S7H), in agreement with phalloidin staining that showed a dense and compact morphology of Atoh8-KD-derived cells compared to the more elongated and less clustered Ctrl-derived ones (Fig. 6J). These morphological differences prompted us to investigate whether Atoh8 controls phenotypic plasticity by constraining changes in the mesenchymal/epithelial status of the cells. In line with this hypothesis, we observed a significant increase of the epithelial transcripts *Cdh1, Epcam* and *Krt18* in Atoh8-KD-derived cells but no concomitant decrease in the expression of the mesenchymal markers *Vim* or *Snail* (Fig. 6I), potentially indicative of a partial EMT state, as reported in skin and mammary tumor models as well as in human head and neck cancers (*47, 48*). We next showed that Cdh1 is significantly induced at the protein level in three independent Atoh8-KD-derived cell lines (Fig. 6K and S7I) and that this constitutes an early event detected as early as 3 days after the induction of MT (Fig. 6L). Because tumorigenic populations encompassing partial EMT state were found to be highly plastic and aggressive, we assessed the tumorigenic potential of Ctrl- and Atoh8-KD-derived lines both *in vitro* and *in vivo.* Atoh8-KD-derived lines were significantly more prone to grow in non-adherent conditions (Fig. S7J-M). To rule out the possibility of a stochastic clonal expansion, the experiment was reproduced with three independent lines, obtaining similar results (Fig. S7N-O). Injection into immunocompromised mice showed an increased growth of Atoh8-KD-derived tumors and a significant reduction of the overall mice survival (Fig. 6M-O). Collectively, the data indicate that Atoh8 constrains the acquisition of malignant properties but also the emergence of phenotypically plastic and aggressive tumor cells.

### Definition of the Atoh8 genome-wide binding and function in cellular plasticity

Due to the broad ability of the bHLH TF Atoh8 to constrain cellular plasticity during reprogramming, trandifferentiation and transformation, and assuming that TF binding determines function, we assessed its genomic distribution after ectopic expression of an AM-tagged version of Atoh8 in MEF (Fig. S8A). Chromatin immunoprecipitation (ChIP) followed by sequencing led to the identification of 1826 peaks, principally distributed in upstream/downstream gene regions and introns (Fig. 7A-B). The ChIP-seq signal specificity was confirmed with the mock sample (AM-tag vector) which lacked signals at the corresponding peaks (Fig. 7C). Motif analysis showed that Atoh8 binds preferentially the CAGCTG motif (E-box), even if the TGACTC motif (AP-1) was also enriched (Fig. 7D).

**Figure 7:**
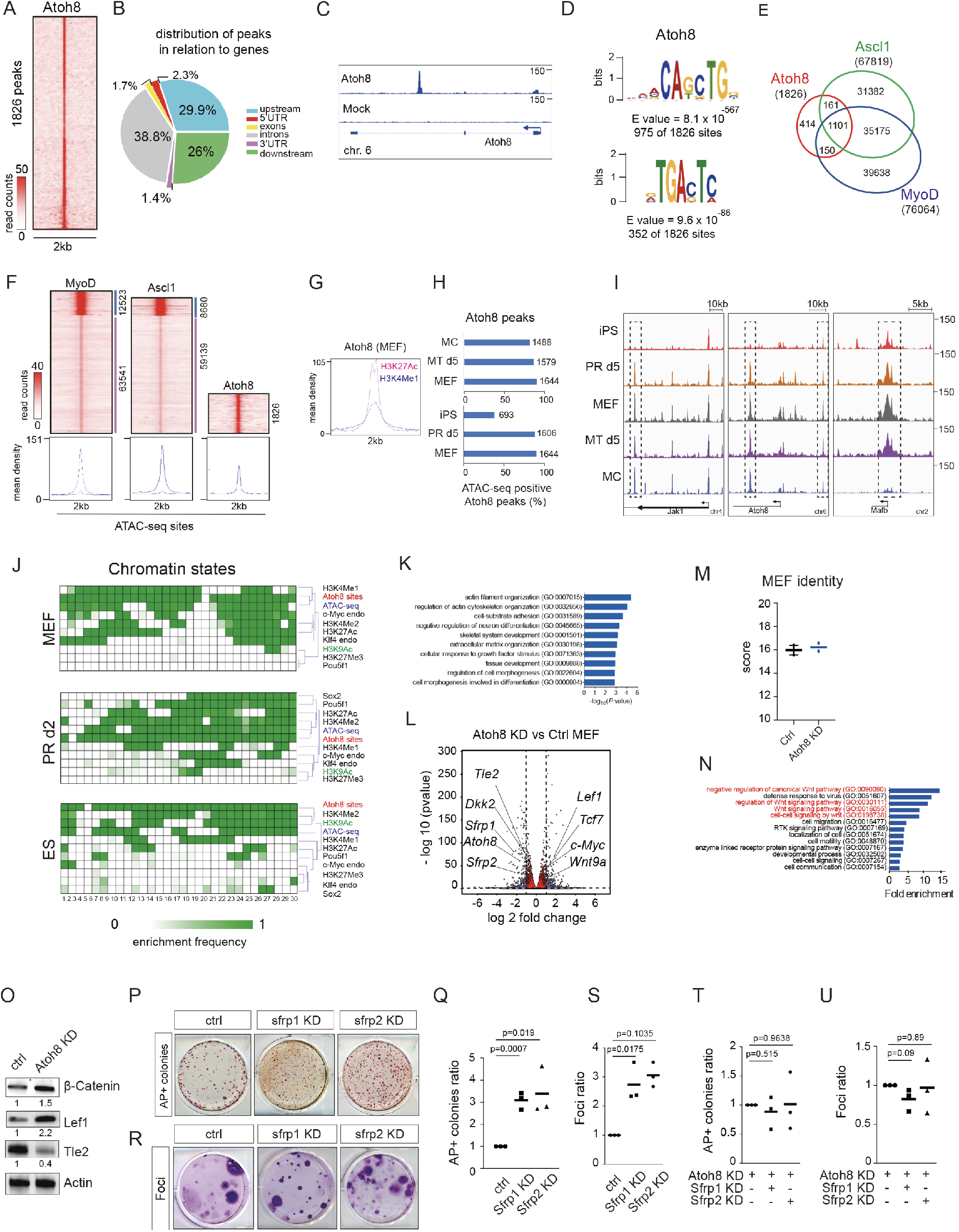
Atoh8 constrains cellular plasticity by fine-tuning WNT signalling activity. (A) Heat map displaying Atoh8 read counts +/-1kb around merged peak summits. (B) Genomic distribution of Atoh8-specific peaks revealing its preference to intergenic/intronic regions. (C) Genome browser track showing Atoh8 binding within its own intronic region. (D) Most enriched DNA-binding motifs associated to Atoh8 issued from a *de novo* motif analysis (MEME). (E) Venn diagram displaying the fraction of common peaks between Atoh8 (this study), Ascl1 and MyoD (*37*). (F) Heat map displaying the enrichment of MyoD, Ascl1 or Atoh8 within chromatin accessible sites (ATAC-seq). Notice that, in contrast to Atoh8 binding, only a small fraction of MyoD and Ascl1 sites are retrieved within highly enriched ATAC-seq sites. (G) mean read counts enrichment density associated to the active histone modification marks H3K27Ac and H3K4Me1 within the Atoh8-specific sites. (H) Fraction of Atoh8 sites retrieved within open chromatin regions (ATAC-seq) assessed during MEF reprogramming towards pluripotent stem (iPS) cells or transformation into malignant cells (MC). (I) example of open chromatin sites at different stages during MEFs reprogramming and transformation. (J) Atoh8-centered Chromatin state analysis revealing its binding co-occurrence with chromatin epigenetic marks and pluripotency TF (*6, 68*). (K) GO analysis of the 724 genes located in the vicinity (100kb) of an Atoh8 ChIP-seq peak. (L) Volcano plot comparing the transcriptomes of control (Ctrl) and Atoh8 KD MEF. Each dot corresponds to a transcript. Blue dots present a log2 fold change >0.8 or <-0.8, and adjusted p-value <0.00001. (M) Effect of Atoh8 KD on MEF identity score. SSgsea analysis based on 395 genes of MEF identity depicted in (*9*). (N) Graph depicting fold enrichment of statistical overrepresentation analysis. The Pantherdb tool was used to detect overrepresented GO terms within the genes differentially expressed in control and Atoh8 KD MEF. A Fisher’s exact two-sided test was used to calculate p-values. (O) Western blot of β-catenin, Lef1 and Tle2 in control and Atoh8 KD settings. (P) Picture representing AP+ colonies at day 15 of PR in Control (Ctrl), Sfrp1 KD and Sfrp2 KD settings, representative of two independent experiments. (Q) Colony counting. Data are the mean ± s.d. (n=2 independent experiments). (R) Picture representing Cresyl-violet foci formation colonies at day 30 of MT in Control (Ctrl), Sfrp1 KD and Sfrp2 KD settings, representative of three independent experiments. (S) Colony counting. Data are the mean ± s.d. (n=3 independent experiments). Student’s t-test was used, and two-sided p-values are indicated. (T) Colony counting of AP+ colonies at day 15 of PR in Atoh8 KD, Atoh8+Sfrp1 KD and Atoh8+Sfrp2 KD settings. Data are the mean ± s.d. (n=3 independent experiments). Student’s t-test was used, and two-sided p-values are indicated. (U) Counting of Cresyl-violet foci formation colonies at day 30 of MT in Atoh8 KD, Atoh8+Sfrp1 KD and Atoh8+Sfrp2 KD settings. Data are the mean ± s.d. (n=2 independent experiments).

Wernig and colleagues recently demonstrated that the bHLH class A pioneer TFs Ascl1 and MyoD, that trigger respectively the transdifferentiation of MEF towards neuronal or muscular fates, bind the same CAGCTG E-box motif in MEF (*37*). By integrating the corresponding datasets, we found first that a large fraction of the Atoh8 peaks (60%) become occupied by both Ascl1 and MyoD during MEF transdifferentiation (Fig. 7E). However, the general features of Atoh8, Ascl1 and MyoD distribution were different. In contrast to Ascl1 and MyoD that mainly bind inaccessible regions as pioneer factors, Atoh8 was found principally in accessible (ATAC-seq positive) enhancer regions enriched in H3K27Ac and H3K4Me1 signals, distant from TSS in MEF (*6*) (Fig. 7F-G and S8B).

We next assessed how the chromatin accessibility of the Atoh8-bound regions behave during PR and MT. We found that they remained largely accessible (ATAC-seq positive) after 5 days of PR or MT and in malignant cells but a significant loss of ATAC signal was observed in iPS cells (Fig. 7H). As examples, the chromatin accessibility of the Atoh8-bound regions in the Atoh8 gene is progressively reduced during both PR and MT, in adequation with its downregulation in iPS and malignant cells (Fig. 4I). To better dissect the dynamic of the Atoh8-bound peaks during PR, we conducted a chromatin combinatorial states analysis combining ChIP-seq datasets for chromatin accessibility (ATAC-seq), histone marks (H3K4Me1, H3K27Ac, H3K4Me2, H3K27Me3, H3K9Ac) and TFs (Oct4-Pou5f1, Sox2, Klf4 and c-Myc) in MEF, after 48 hrs of OSKM expression and in embryonic stem (ES) cells (*6, 49*). In line with our results, we first observed that the chromatin accessibility of the Atoh8-bound regions was maintained after 2 days of PR but significantly reduced in ES cells (ATAC). Moreover, those regions progressively lost the H3K27Ac and H3K4Me1 enhancer marks while concomitantly gained H3K9Ac signal in ES cells (Fig. 7J). Of note, a significant proportion of Atoh8 peaks was also found to be co-occupied by c-Myc in MEF, but this fraction dropped significantly after 2 days of PR and in ES cells, indicating a gradual relocation of c-Myc, paralleled by a transient binding of Oct4 and, to a lesser extent, Sox2, observed at day 2. Altogether, these data indicate highly dynamic epigenetic reconfigurations of the Atoh8-bound regions during reprogramming and transformation.

Because Atoh8 is mainly bound to enhancer regions in MEF, we focused next on the genes that it might regulate. Pantherdb overrepresentation tests conducted on the 724 genes located in the vicinity of the Atoh8-bound peaks (<100 kbs) highlighted significant enrichment for genes related to “cell adhesion” (Col1a1, Lamb1) (*37*), but also more broadly with “developmental process” (Runx1, Runx3, Notch2, Tead1) and “neuron differentiation” (Hes1, Id1, Sema6a) (Fig. 7K). We also noticed a direct binding of Atoh8 on its own locus (Fig. 7C), suggesting the existence of an auto-regulatory loop (*50*), and to a significant number of bHLH promoters including Id1, Id2, Id3, Hes1, Bhlhe41, Twist2, Max, Tfap4 and Arnt2. Of note, a large number of WNT signalling pathway genes such as Wnt9a, Wnt10a, Tle3, Tle6, Kremen1, Axin2 and Wisp2, were bound by Atoh8. To refine the gene regulatory network controlled by Atoh8, we conducted RNA-seq on MEF control (Ctrl) or knockdown (Atoh8 KD) for Atoh8 for 5 days. Bio-informatic analyses indicated the deregulation of 503 genes (fold change log2 > 0.8 et < – 0.8, adjusted pvalue <10-^5^, 229 up, 274 down) (Fig. 7L). We found that 43 deregulated genes also harbored Atoh8 ChIP-seq peaks, such as the Mafb and Jak1 genes depicted in Figure 7I. Interestingly, among those 43 genes, 23 were downregulated and 20 upregulated, suggesting that Atoh8 may exert both activating and repressing functions (Fig. S8C). We noticed that Atoh8 KD led to a significant induction of c-Myc at both transcript and protein levels (Fig. 7L and S8D). On the contrary, c-Myc was found to bind to Atoh8 promoter and to repress its expression (Fig. S8E-F), indicating the existence of a Atoh8/c-Myc negative feedback loop. Importantly, Atoh8 depletion had no significant effect on MEF identity score, suggesting that its primary function is not to safeguard cellular identity (Fig. 7M). Pantherdb overrepresentation assays conducted on the genes deregulated by Atoh8 KD indicated a putative link with the WNT signalling pathway, with significant downregulation of the WNT inhibitors *Sfrp1, Sfrp2, Dkk2* and *Tle2* and concomitant upregulation of the WNT effectors *Wnt9a, Tcf1* and *Lef7* (Fig. 7L and N). These results prompted us to investigate whether Atoh8 controls WNT signalling activity. Tle2 downregulation, concomitant with Lef1 and active β-catenin upregulation, were observed following Atoh8 knockdown in MEF (Fig. 7O). Next, to evaluate whether Atoh8 might constrain cellular plasticity by fine-tuning WNT *via* Sfrps levels, we assessed whether Sfrp1 and Sfrp2 knockdown mimicks the effects of Atoh8 depletion on PR and MT. Sfrp1 and Sfrp2 KD in MEF (>50% knockdown efficiency) (Fig. S8G) led to a 2.9- and 2.3-fold increase of the number of AP+ colonies (Fig. 7P-Q), and 2.7- and 3-fold more immortalized foci (Fig. 7R-S). Importantly, by devising a combinatorial RNAi strategy, we found that the simultaneous suppression of Atoh8 and Sfrp1, or Atoh8 and Sfrp2, impaired PR (Fig. 7T) and MT (Fig. 7U), in a similar range as the suppression of Atoh8 alone, indicating that Atoh8 depletion effect on PR and MT is mediated by a unique axis involving Sfrps. Altogether our results reveal how Atoh8 constrains cellular plasticity by binding to specific enhancers and fine-tuning WNT signalling activity.

## Discussion

Despite the critical function of cellular identity loss and cellular plasticity acquisition in both induced pluripotency and cancer biology, the associated molecular circuitries and their degree of analogy remain poorly defined, mainly because of lacking genetic tools allowing sophisticated comparative analyses. We exploited in this study “repro-transformable” mice to compare the early molecular and cellular responses to the induction of reprogramming and transformation. First, we revealed that OSKM prevent DNA damage induced by various oncogenic factors including c-Myc and K-Ras in MEF. This effect is complementary to their known protective role on ageing hallmarks (*51*) and, in the context of the link between OSKM, ageing, senescence and cancer development (*51–54*), it will be highly relevant to assess whether this phenomenon also occurs *in vivo* during tumorigenic processes.

PR and MT are poorly efficient and few cells engage in the processes in the first days. Characterization of bulk populations has provided some insights, but as most cells fail to generate iPS/malignant cells, those analyses are necessarily biased toward measurement of unproductive reprogramming/transforming events. Recent single-cell reports on PR infer the trajectories and the potential of single cells based on transcriptome similarities (*8, 10,11*). Here, in a complementary approach, we attempted to identify somatic markers allowing the isolation of early populations of cells that functionally gained plasticity. In contrast to previous reports on PR (*3, 30*), we exploited the sequential downregulation of somatic markers to comparatively trace the route maps of PR and MT. By combining functional assays with epigenomic and transcriptomic analyses of cellular intermediates (CI), we revealed that similar transcriptomic and epigenomic modifications occur in the early phase of PR and MT, reinforcing the concept of the analogy between both processes. We detected in particular an unexpected and transient induction of immune- and inflammation-related genes early in PRP and MTP cells. Even if additional efforts will be required to examine the beneficial effects of single, or combination of such molecules on PR and MT (*52, 53, 55*), it is relevant to note that similar increased expression was recently discovered as a feature of transitional cellular states in melanoma and lung cancer (*56, 57*). We also showed that cellular identity loss and cellular plasticity acquisition are uncoupled during PR and MT. Indeed, the PRP and MTP CI harbor a strong reprogramming/transforming potential but no significant downregulation of their MEF identity score yet. This finding appeared to us conceptually important for regenerative and cancer biology.

We identified the bHLH TF Atoh8, initially described as a key factor for neurodevelopment (*58*), as a broad-range regulator of iPS cells generation, transdifferentiation and malignant transformation. Atoh8 function during PR was still under debate with contrasting pro- and anti-reprogramming roles described so far (*46, 59*). Based on our findings, we propose that Atoh8 does not safeguard cellular identity but rather prevents the acquisition of cellular plasticity. In line with this view, Atoh8 depletion led to a rapid gain of phenotypic plasticity during malignant transformation, leading to the establishment of highly aggressive population of cells expressing both epithelial and mesenchymal markers. It will be of great importance to analyze these populations at the single-cell level to evaluate whether cells co-express these markers, as recently reported in various cancers (*47, 48*). Defining the gene regulatory network controlled by Atoh8 in MEF led to a similar view. Atoh8 genome-wide binding revealed a restricted binding to enhancer regions that are enriched for endogenous c-Myc in MEF. However, during reprogramming and transdifferentiation, these regions are dynamically regulated with binding of Oct4/Sox2 during PR and Ascl1/MyoD during transdifferentiation. Atoh8 depletion does not impact significantly cellular identity but rather fine-tunes the expression of several Wnt signalling members, well described to promote plasticity in regeneration and cancer (*60, 61*). Because Wnt signalling has been found to exert stage-specific functions during PR (*62, 63*), we propose that c-Myc-mediated downregulation of Atoh8 in a subset of cells might control PR/MT by contributing to activate Wnt signalling.

By deciphering the early molecular routes of reprogramming and transformation, by uncoupling cellular identity loss and cellular plasticity acquisition, and by revealing antagonistic functions for bHLH TF networks, our work provides a conceptual framework that opens fascinating perspectives for regenerative medicine and cancer biology.

## Methods

### Mice and MEFs

R26^rtTA^;Col1a1^4F2A^ (*64*), LSL-K-ras^G12D^ (*65*), R26-CRE^ERT2^, Oct4-EGFP and Bcl11b-tdTomato (*33*) mice were housed under standard conditions and bred in accordance with french national guidelines. Genotyping was carried out on genomic DNA derived from adult and embryonic tails using the DirectPCR Lysis Reagent (102-T, Viagen Biotech) and EconoTaq Plus Green 2X Master Mix (Lucigen). Primers used are listed in Table 1.

MEFs were isolated from E13.5 embryos after removal of the head and internal organs. The remaining tissues were physically dissociated and incubated in trypsin at 37°C for 10 min after which cells were resuspended in MEF medium.

### Teratoma

Teratoma formation assays were performed by injecting 1×10^6^ iPS cells into the testes of 7-week-old severe combined immunodeficient (SCID) mice (CB17/SCID, Charles River). After 3-4 weeks, the mice were euthanized and lesions were surgically removed and fixed in 4% paraformaldehyde for sectioning and hematoxylin-eosin staining.

### Plasmids and constructs

pMXS-Oct4, pMXS-Sox2, pMXS-Klf4, pMXS-Myc, pLKO.1 and pWPXLd plasmids were purchased from Addgene. shRNAs against Trp53, FosL1, Atoh8, Sfrp1 and Sfrp2 were designed using the MISSION shRNA library from Sigma-Aldrich and ligated using the Rapid DNA ligation kit (Sigma-Aldrich) into the pLKO.1 vector digested with AgeI and EcoRI. shRNA sequences are listed in Table 1. Atoh8 cDNA was amplified from MEFs and cloned into the pWPXLd expression vector at BamH1 restriction site (AM-Tag is added at the C-terminal). Single guide RNA targeting Atoh8 (designed with CRISPOR program) were cloned into the lentiCRISPRv2 plasmid at a BsmBI restriction site. Single guide RNA sequences are listed in Table 1. pWPIR Hras G12V and cyclinE plasmids were kindly supplied by A. Puisieux’s lab. Tet-O-FUW-Brn2, Tet-O-FUW-Ascl1, Tet-O-FUW-Myt1l and FUdeltaGW-rtTA plasmids were purchased from Addgene.

### Cell culture and viral production

MEF medium consists of DMEM supplemented with 10% fetal bovine serum (FBS), 100 U/mL penicillin / streptomycin (PS), 1 mM sodium pyruvate, 2 mM L-glutamine, 0.1 mM Non Essential Amino Acids (NEAA) and 0.1 mM β-mercaptoethanol.

pMXs-based retroviral vectors were generated with Plat-E cells (a retroviral packaging cell line constitutively expressing *gag, pol* and *env* genes). Briefly, calcium phosphate transfection of the vectors was performed with the CalPhos Mammalian Transfection kit (Ozyme) in 10-cm dishes. Medium was changed with 10 mL of MEF medium after 7 h of incubation. The lentivirus-containing supernatants were collected 48 h later and stored at −80 °C. 293FT cells, grown in MEF medium, were used to produce lentiviral particles. The vectors were transfected along with plasmids encoding the envelope G glycoprotein of the vesicular stomatitis virus (VSV-G) and Gag-Pol.

### Pluripotent reprogramming

For Dox-induced PR, reprogrammable R26^rtTA^;Col1a1^4F2A^; Oct4-EGFP MEFs within three passages were plated in six-well plates at 80,000-100,000 cells per well in MEF medium. The following day, cells were infected overnight with shRNA-carrying lentiviral stocks in the presence of 8 μg/ml polybrene, and medium was then replaced by fresh medium with 2 μg/ml Dox. MEFs were reseeded 72 h after infection on 0.1% gelatin coated plates in iPSC medium (DMEM containing 15% KnockOut Serum Replacement, 1,000 U/mL leukemia inhibitory factor, 100 U/mL PS, 1 mM sodium pyruvate, 2 mM L-glutamine, 0.1 mM NEAA and 0.1 mM β-mercaptoethanol, at equal densities for each condition to normalize potential effect of differential MEF proliferation on reprogramming efficiency. Several densities were tested (15,000-68,000 cells per cm^2^). Every day, medium was either replaced by or supplemented with Dox-containing fresh medium. Once iPS colonies were macroscopically visible, OCT4-EGFP*+* colonies were counted under an Axiovert 200 M microscope, and alkaline phosphatase (AP) staining was performed using the Leukocyte Alkaline Phosphatase kit (Sigma-Aldrich). Alternatively, MEFs were coinfected with OSKM retroviral vectors 48 h after lentiviral infections and cultured identically thereafter.

For human pluripotent reprogramming, human dermal fibroblasts (HDF, Sigma) were cultivated in MEF medium and were infected with lentiviral sgRNA particules in the presence of 8μg/mL polybrene and medium was replaced by fresh medium the following day. Two days after sgRNA infection, HDFs were infected with OSKM Sendai particles (CytoTune™-iPS 2.0 Sendai Reprogramming Kit, Life technologies) and medium was replaced by fresh medium the following day and every other days until day 9. After 9 days, cells were splitted onto vitronectin and medium was changed to mTESR medium (Stem cell technologies).After approximately 26 days, colonies were SSEA4 live-stained (GloLIVE Human Pluripotent Stem Cell Live Cell Imaging Kit, ReD) and counted under an Axiovert 200 M microscope. Alternatively, alkaline phosphatase (AP) staining was performed with the Leukocyte Alkaline Phosphatase kit (Sigma-Aldrich).

### Malignant transformation

For MT, the LSL-K-ras^G12D^;R26-CRE^ERT2^ MEF were similarly infected overnight with shRNA-carrying lentiviral stocks in the presence of 8 μg/mL polybrene. 48 h later the cells were coinfected overnight with *shTrp53-* and *Myc*-carrying viruses concomitantly with 4-hydroxitamoxifen treatment (1 μM) to induce K-ras^G12D^ expression. Alternatively, the coinfection of *shTrp53-, Myc-* and Hras^G12V^-carrying viruses was used in WT MEF to initiate MT. MEF were reseeded 48 h post-infection in six-well plates at low density (500, 1,000 or 2,000 cells per well) in focus medium (MEF medium with 5% FBS) for the foci formation assay. Medium was then changed twice a week. After several passages of the cells derived from MT, soft agar assays were performed. Transformed cells were plated on an agarose-containing MEF medium layer at a density of 25,000-50,000 cells per six-well plate. Foci and soft agar colonies were stained 25-30 days later with a 0.5% cresyl violet solution in 20% methanol.

### Xenografts

3×10^6^ immortalized cells were prepared in 100μl PBS supplemented with 100μl matrigel and injected subcutaneously into immunocompromised SCID mice (N=6 for each group). The volume of the tumor is then measured every 3 days until day 16.

### CAM assay

2.5×10^6^ immortalized cells were inoculated on the chorioallantoic membrane (CAM) in the egg of chick embryos at E11 where they form a primary tumor. The size of the tumor was evaluated after 7 days. The number of replicates in indicated in the figure legend.

### MEF to neurons transdifferentiation

WT MEF were co-infected with FUdeltaGW-rtTA and sgRNA (control or targeting Atoh8) lentiviral plasmids at day −2 in the presence of 8 μg/ml polybrene. At day 0, cells were coinfected with Tet-O-FUW-Brn2, -Ascl1 and -Myt1l lentiviral plasmids. The day after, the medium was replaced by fresh MEF medium supplemented with 2 μg/ml Dox. At day 3, the medium was replaced by fresh N3 medium consisting in DMEM-F12, 100U/ml penicillin / streptomycin (PS), 2.5μg/ml insulin, 50μg/ml apo-transferrin, 86.5μg/ml sodium selenite, 6.4ng/ml progesterone, 16μg/ml putrescine supplemented with 2 μg/ml Dox. The medium was changed daily until day7-8.

### Immunofluorescence

Cells were fixed with 4% paraformaldehyde for 10 min at room temperature (RT°C), washed 3 times with PBS, permeabilized with 0.1% Triton X-100 for 30 min at RT°C and blocked with 1% bovine serum albumin (BSA) for 1 h. After incubation with primary antibodies against NANOG (Reprocell, RCAB001P), SSEA1 (Santa Cruz Biotechnology, sc-101462), OCT4 (sc-5279, Santa Cruz), SOX2 (ab97959, Abcam), MAP2 (Sigma Aldrich, M4403), and phosphoHistone H2A.X (Cell Signaling Technology, 2577) overnight at 4°C, cells were washed 3 times with PBS and incubated with fluorophore-labeled appropriate secondary antibodies (Life Technologies). Phalloidin staining was performed using GFP-coupled Phalloidin-Atto 488 (49409, Sigma-Aldrich). Live SSEA4 immunostaining was carried out with the GloLIVE Human Pluripotent Stem Cell Live Cell Imaging Kit (SC023B, R&D).

### RNA extraction and RT-qPCR

Total RNAs were extracted using Trizol reagent and 1 μg of RNA was reverse-transcribed with the RevertAid H Minus First Strand cDNA Synthesis kit (Life Technologies). qPCR was performed with the LightCycler 480 SYBR Green I Master mix (Roche) on the LightCycler 96 machine (Roche). *Gapdh* and *Rplp0* were used as housekeeping genes. qPCR primers are listed in Table 1.

### Chromatin immunoprecipitation

MEFs were infected with lentiviral particles carrying AM-tagged Atoh8. After 3 days, DNA was extracted, precipitated and purified using the Tag-ChIP-IT kit (53022, Actif Motif). qPCR were performed as described above for ChIP-qPCR.

### Protein extraction and Western blot

Cells were harvested in RIPA buffer (150 mM NaCl, 1% Triton, 0.5% deoxycholate, 0.1% SDS and 50 mM Tris, pH 8.0) supplemented with protease inhibitors and phosphatase inhibitors. After 30 min on ice, lysis by sonication, and centrifugation for 10 min at 15,000*g*, supernatants were collected, proteins were denatured 10 min at 95 °C in Laemmli sample buffer, separated on 4-15% polyacrylamide gel, and transferred onto a nitrocellulose membrane. The membrane was blocked with 5% milk in TBST (Tris-buffered saline, 0.1% Tween 20) for 1 h, incubated with primary antibody at 4°C overnight and secondary antibodies for 1 h at RT°C. Antigens were detected using ECL reagents. The following antibodies were used: mouse anti-OCT4 (sc-5279, Santa Cruz, 1:1000), rabbit anti-SOX2 (ab97959, Abcam, 1:1000, mouse anti-c-MYC (sc-42, Santa-Cruz, 1:200), rabbit-anti ATOH8 (PA5-20710, Termofisher, 1:1000), rabbit anti-ID4 (BCH-9/82-12, BioCheck, 1:1000), mouse anti-TWIST2 (HOO7581-M01, Abnova, 1:250), rabbit anti-NANOG (RCAB002P, Reprocell, 1:1000), mouse anti-SSEA1 (sc-101462, Santa Cruz, 1:1000), mouse anti-CDH1 (610181, BD, 1:1000), rabbit anti-SNAIL (C15D3, Cell signaling, 1:1000), rabbit anti-VIM (R28, Cell signaling, 1:1000), mouse anti-TWIST1 (ab50887, Abcam, 1:250), goat anti-hSOX2 (AF2018, R&D, 1:1000), rabbit anti-hNANOG (3580, Cell signaling, 1:1000), mouse anti-ACTIVE β-CATENIN (8E7, Millipore, 1:1000), mouse anti-TLE2 (sc-374226, Santa-Cruz, 1:500), rabbit anti-LEF1 (2230, Cell signaling, 1:1000), rabbit anti-GAPDH (sc-25778, Santa-Cruz, 1:4000),rat anti-BCL11B (Abcam, ab18465, 1:500), horse radish peroxidase (HRP)-conjugated anti-ACTIN (Sigma Aldrich, A3854, 1:10,000).

### FACS

The following antibody was used: anti-mouse CD90.2 (Thy-1.2) APC (eBioscience, 17-0902), CD73-AF488 (BD Biosciences, 561545), CD49d-PE (BioLegend, 103705). Analysis was performed on a BD LSRFortessa. Sorting was performed on a BD FACSAria. Apoptosis was measured using the FITC Annexin V/Dead Cell Apoptosis Kit (Invitrogen, V13242). For cell cycle analysis, the cells were fixed in ethanol 70% and stained with 40 μg/mL propidium iodide supplemented with 2 mg/mL RNase.

### NGS analyses

RNA quality was analysed using a Bioanalyser (Agilent). Libraries were constructed and sequenced on an illumina Hiseq 2000 by the cancer genomics platform on site. ATAC-seq, and ChIP-seq data were generated by the Active Motif company. Atoh8 ChIP-sequencing and ATAC-seq datasets were aligned to the mouse reference genome assembly mm10 using Bowtie 2.1.0 under by default parameters. Peak calling was performed using MACS 2.1.1. For experiments with replicates, BED files obtained after alignment were concatenated prior MACS peak calling processing. Atoh8-specific sites were obtained by subtracting binding sites observed within the AM-tag processed control dataset (BEDTools 2.29.2). Enrichment heat maps and mean density plots were obtained with seqMINER (*66*). De novo motif analysis has been performed with MEME-ChIP (*67*). Read counts enrichment signals were visualized with IGV genome browser. Atoh8-centered chromatin state analysis has been performed by intersecting Atoh8-specific binding sites with those associated to public data (*6, 68*) then inferring co-occuring events with ChromHMM (*69*).

### Statistical analyses

All data are reported as mean ± SEM. Statistical analyses were performed using GraphPad Prism software. Student t tests were used for paired comparisons and one-way ANOVA followed by a Tukey’s post hoc test was used for multiple comparisons. P-values are indicated on each graph.

## Data availability

NGS data were deposited on GEO (record number series GSE137050 and GSE143594, secure token for reviewers access yvmpscquvhybzgl and ejctmmwcnfivrmt).

## Acknowledgements

We are grateful to the Laboratoire des modèles tumoraux (LMT) and A. Lalande for technical assistance.

## Author Contributions

AH and GF performed most of the experiments from figures 1–7. NR, JS and JP performed bio-informatic analyses. BG performed *in vivo* work. FL, AH and GF designed experiments and wrote the manuscript. FL designed and supervised the study.

