## Supplementary material for "The comprehensive roadmaps of reprogramming and transformation unveiled antagonistic roles for bHLH transcription factors in the control of cellular plasticity"

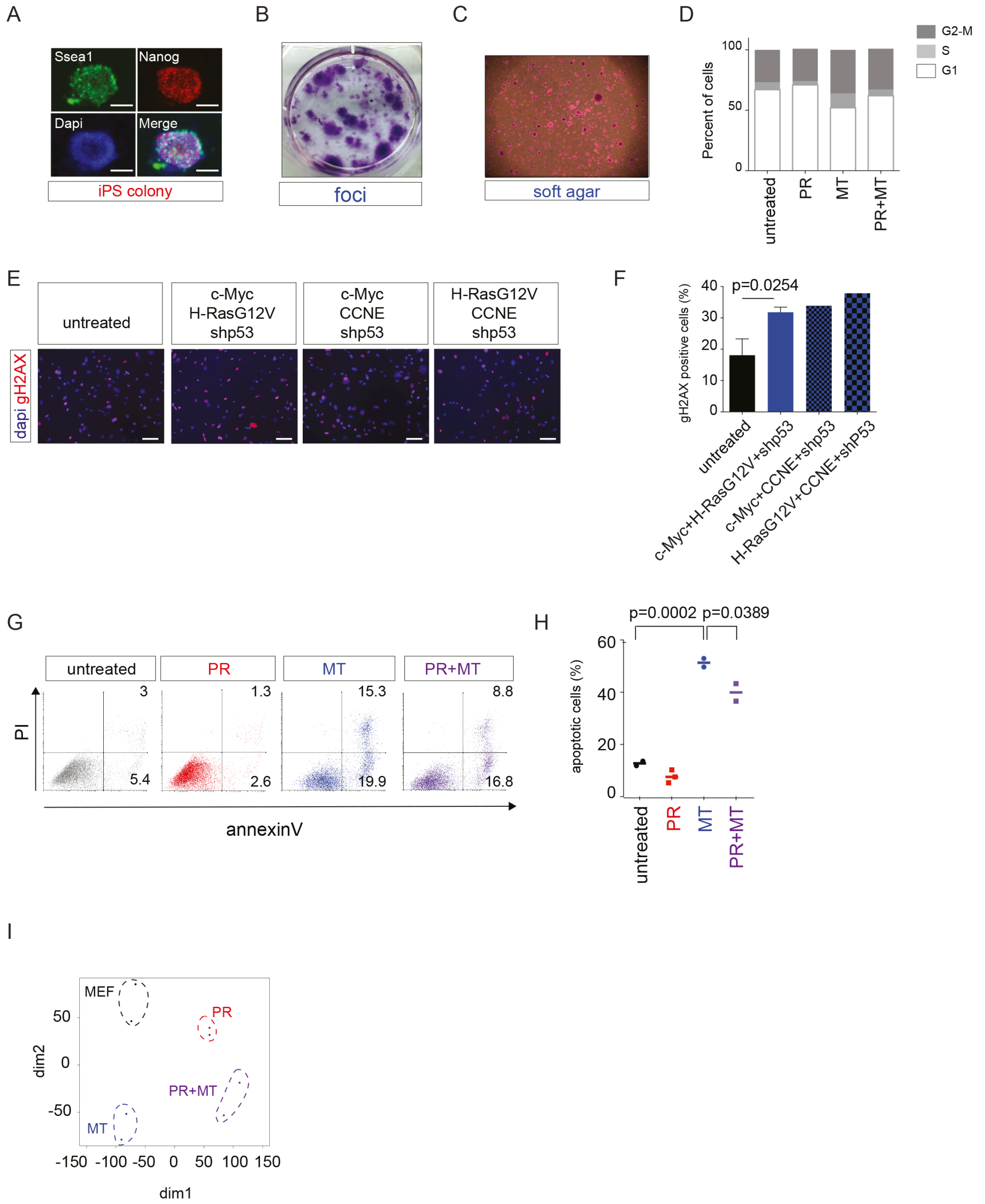


**Supplementary figure 1.** (A) Immunofluorescent staining of PR-induced iPS cells for Ssea1 and Nanog. Scale bar: 100 µm. (B) Foci from MT-induced malignant cells colored with cresyl-violet. (C) Soft agar colonies derived from MT-induced malignant cells colored with cresyl-violet. (D) Cell cycle analyzed by FACS in control MEF and after 5 days of PR, MT and PR+MT. (E) Immunofluorescent staining of MT-induced cells for γH2AX after 3 days with different oncogenic cocktails compared to control MEF. CCNE: Cyclin E ; shp53: shRNA targeting p53. (F) Counting of γH2AX-positive cells depicted in (E). (G) FACS profile PI/AnnexinV after 3 days of PR, MT and PR+MT compared to control MEF. One representative experiment (from at least two independent experiments). (H) Percentage of total apoptosis depicted in (G). n=3 independent experiments. One-way ANOVA followed by a

Tukey's post hoc test was used. (I) Principal component analysis of normalized gene expression of control cells or cells subjected to 5 days of PR, MT or both programs (PR+MT).

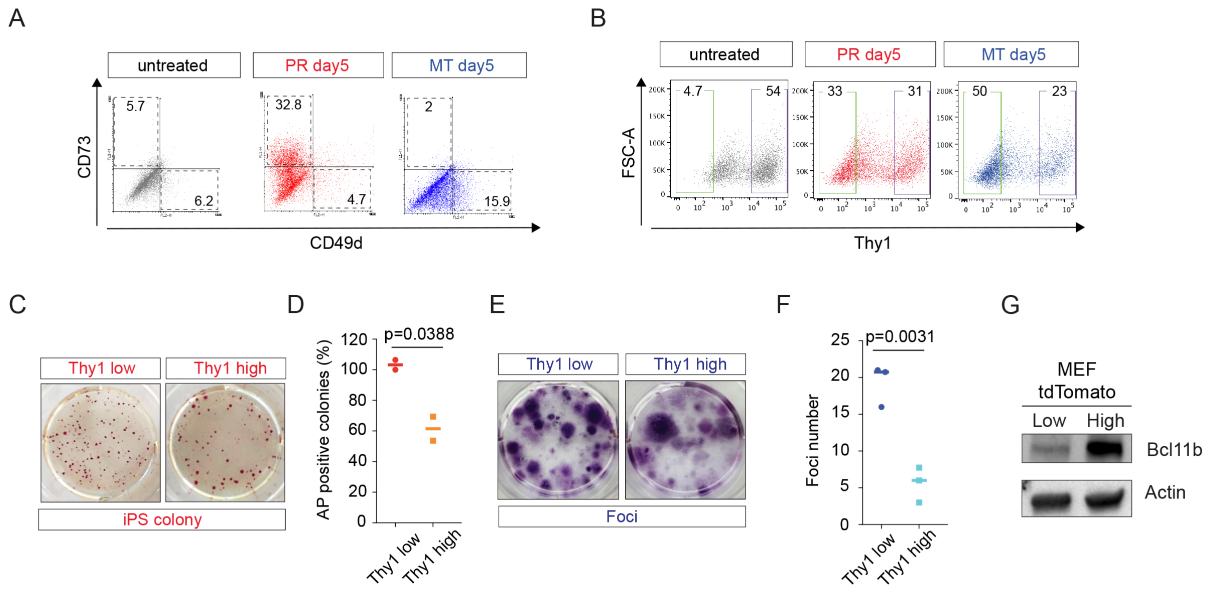


**Supplementary figure 2.** (A) FACS analysis of the CD73 and CD49d after 5 days of PR or MT compared to control MEF. (B) FACS analysis of the somatic marker Thy1 after 5 days of PR or MT compared to control MEF. (C) Alkaline Phosphatase (AP) staining of iPS colonies generated from Thy1$^{low}$ and Thy1$^{high}$ cells FACS-sorted at day 5 of PR. One representative experiment (from two independent experiments). (D) Counting of AP-positive colonies depicted in (C). (E) Foci staining of malignant cells generated from Thy1$^{low}$ and Thy1$^{high}$ cells FACS-sorted at day 5 of MT. One representative experiment (from three independent experiments). (F) Counting of foci depicted in (E). (G) Western blot showing the enrichment of Bcl11b in the tdTomato-high fraction after FACS-cell sorting compared to the tdTomato-low fraction.

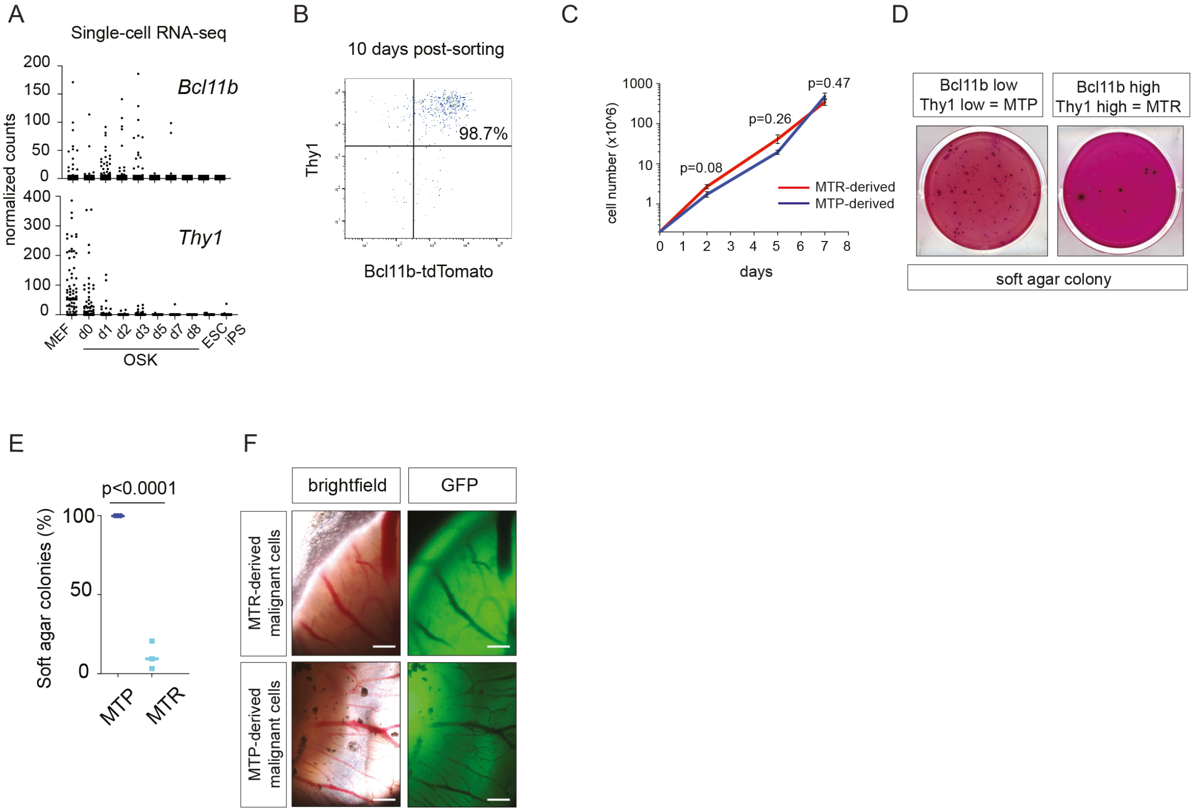


**Supplementary figure 3.** (A) Bcl11b and Thy1 expression at the single cell level during OSK-mediated reprogramming. Data are extracted from (*1*). (B) FACS profile of Bcl11b-tdTomato MEF FACS sorted for high expression of Bcl11b and Thy1 and re-analysed by FACS 10 days later. (C) Growth curves of MTP and MTR-derived cells in 2D. (D) Soft agar assays conducted with independent MTP and MTR-derived cells. (E) Counting of colonies from (D). n=3 independent experiments. Student t test was used. (F) GFP positive cells are present in CAM assays.

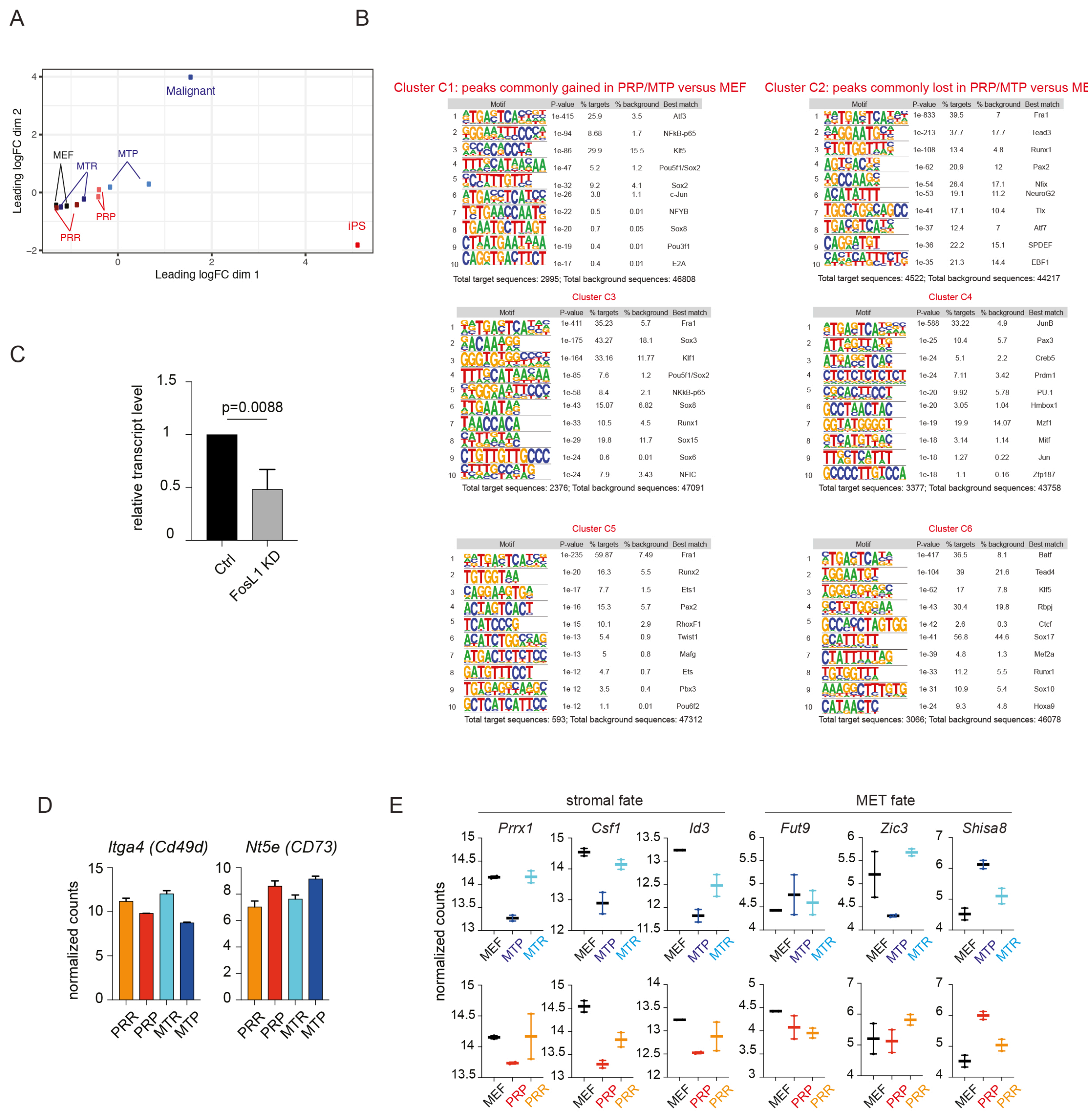


**Supplementary figure 4.** (A) Principal component analysis of ATAC-seq signal of the control cells, the cellular intermediates and the final product of each process (iPS and malignant cells). (B) Top 10 motifs enriched in C1/C2 clusters jointly gained/lost in PRP and MTP, in C3/C4 clusters specifically gained in PRP/MTP and in C5/C6 clusters specifically lost in PRP/MTP. (C) FosL1 knockdown efficiency in MEF. Q-RTPCR of *FosL1* levels 3 days after infection with lentiviral shRNA particles targeting control or FosL1 sequences. Data, normalized to control, are the mean +/- sd of 3 independent experiments. Student T-test was used, and two-sided p-values are indicated. (D) Gene expression levels in MEF and cellular intermediates. Stromal and MET fate genes, as defined in (*2*), were analysed in the RNA-seq datasets from Fig. 4H. (E) *Itga4* and *Nt5e* expression in cellular intermediates during PR and MT. RNA-seq data from fig. 4H are used.

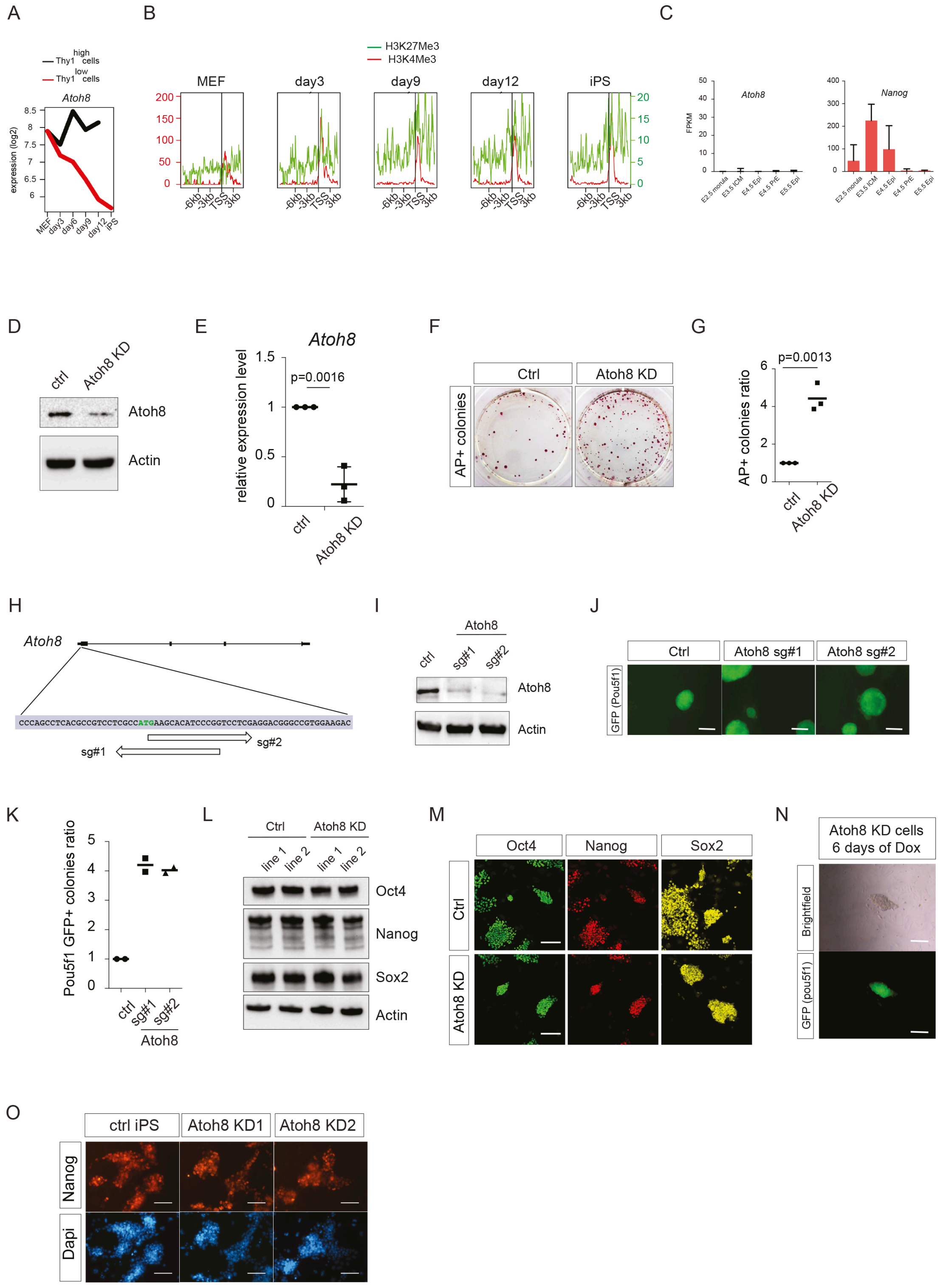


**Supplementary figure 5.** (A) *Atoh8* expression levels in MEF, reprogramming intermediates prone (red - Thy1$^{low}$) and refractory (black - Thy1$^{high}$) populations and iPS cells. Data, extracted from (*3*), are presented as a log2 of transcripts levels in microarray analysis. (B) H3K4Me3 and H3K27Me3 deposition on Atoh8 promoter during iPS cells generation. H3K4m3 (red) and H3K27me3 (green) methylation profile in MEFs and during iPS cells generation, extracted from (*3*). (C) *Atoh8* and *Nanog* expression levels in embryonic stem cells (ESCs) and during pre-implantation development. Data, extracted from (*4*), present transcripts level in FPKM. (D-E) Atoh8 knockdown efficiency in MEF. (D) Western blot of Atoh8 level 3 days after infection with lentiviral shRNA particles targeting control or Atoh8 sequences. (E) Q-RTPCR of *Atoh8* levels in similar settings as (D). Data, normalized to control, are the mean +/- sd of 3 independent experiments. Student T-test was used, and two-sided p-values are indicated. (F) Picture representing AP+ colonies at day 15 of PR in control and Atoh8 knockdown (KD) settings, representative of three independent experiments. (G) AP+ iPSCs colony counting. Data are the mean ± s.d. (n=3 independent experiments). Student's t-test was used, and two-sided p-values are indicated. (H) Scheme depicting the position of the CRISPR/Cas9 guides designed to target the endogenous Atoh8 locus. (I) Efficiency of Atoh8 knock-out (KO) by CRISPR/Cas9 in MEFs. Western blot showing Atoh8 expression level following KO with 2 independent guides. (J) Picture representing Pou5f1-GFP+ iPS colonies at day 15 of PR, representative of two independent experiments. Scale bars=200 µm. (K) Colony counting. Data are the mean ± s.d. (n=2 independent experiments). Student's t-test was used, and two-sided p-values are indicated. (L) Western blot of Oct4, Nanog and Sox2 in independent iPS cell lines obtained in control and Atoh8 KD settings. (M) Immunofluorescence for Oct4, Nanog, and Sox2 in the same lines as (L). Scale bar: 100µm. (N) Brightfield and GFP images showing Pou5f1-GFP+ iPS colonies observed after 6 days of Dox treatment (partial reprogramming). Scale bars=200 µm. (O) Nanog immunofluorescence in *bona fide* and Atoh8-KD iPS independent lines generated a short exposure of 6 days to OSKM expression. Scale bars=500 µm.

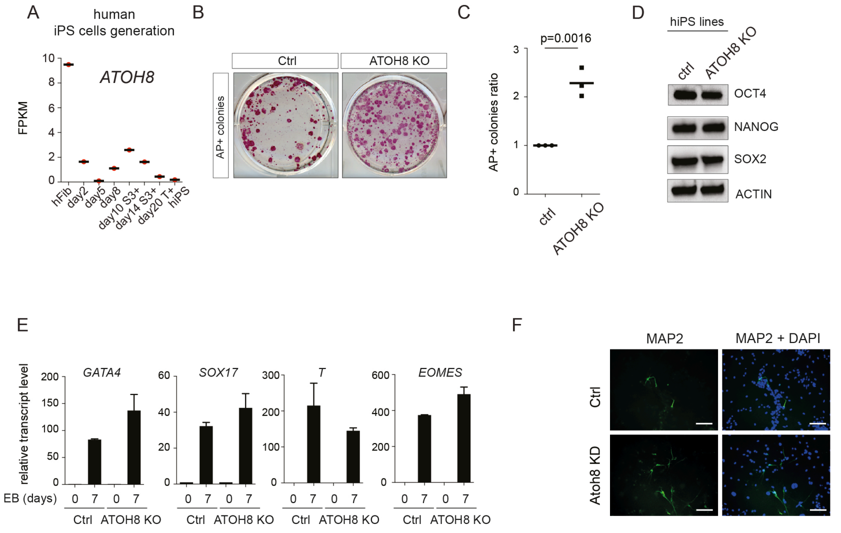


**Supplementary figure 6.** (A) *ATOH8* transcript levels in human dermal fibroblasts, reprogramming intermediates and human iPS cells. Data, extracted from (*5*), present transcripts level in log2 FPKM. (B) Picture representing human AP+ colonies at day 26 of PR in control and ATOH8 KO settings, representative of three independent experiments. (C) Colony counting. Data are the mean ± s.d. (n=3 independent experiments). Student's t-test was used, and two-sided p-values are indicated. (D) Western blot of SOX2, NANOG and OCT4 in hiPS cell lines generated in control and ATOH8 KO background. (E) q-RTPCR showing G*ATA4*, *SOX17, T* and *EOMES* transcript levels during EB formation induced in control and ATOH8 KO iPS cell lines. Data, normalized to day 0 of differentiation, are the mean +/- sd of 3 independent experiments. Student T-test was used, and two-sided p-values are indicated. (F) MAP2 immunostaining performed on iN cells in control (ctrl) and Atoh8 KD settings. Scale bars=100 μm.

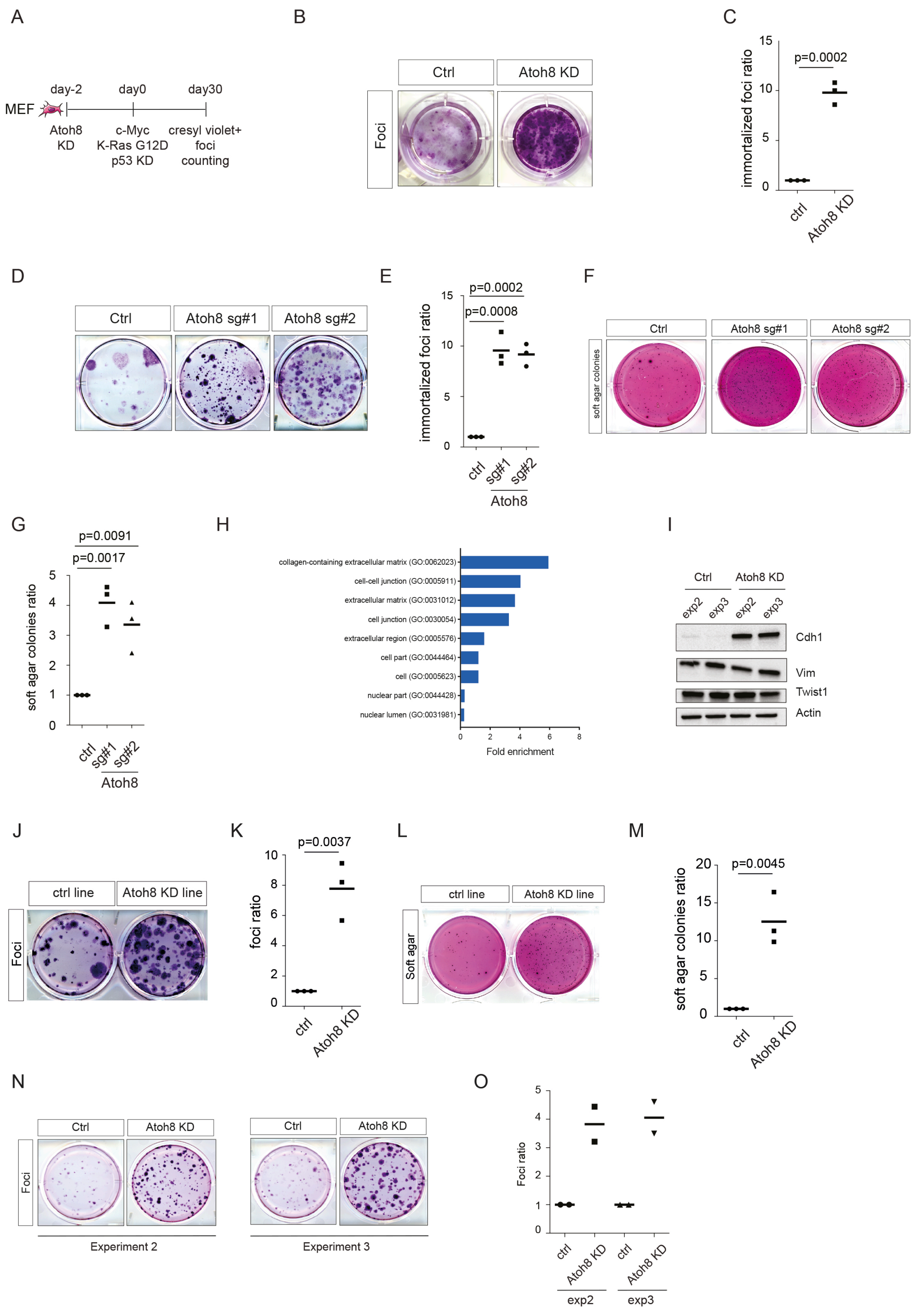


**Supplementary figure 7.** (A) Experimental scheme depicting MEF immortalization. Cells were infected with lentiviral particles targeting control and Atoh8 sequences. 48 hours later, malignant transformation was induced by combining 4-OHT treatment (induces K-RasG12D expression), lentiviral shRNA particles targeting p53 and retroviral particles inducing c-Myc exogenous expression. Immortalized foci were scored at day 30 by Cresyl-violet staining. (B) Picture representing Cresyl-violet foci at day 30 of MEF immortalization in control and Atoh8 KD conditions, representative of three independent experiments. (C) Colony counting. Data are the mean ± s.d. (n=3 independent experiments). Student's t-test was used, and two-sided p-values are indicated. (D) Picture representing Cresyl-violet immortalized foci at day 30 of MEF immortalization upon Atoh8 knock-out using two independent CRISPR/Cas9 guides. (E) Colony counting. Data are the mean ± s.d. (n=3 independent experiments). Student's t-test was used, and two-sided p-values are indicated. (F) Picture representing Cresyl-violet transformed soft-agar colonies at day 30 of malignant transformation following Atoh8 KO with two different CRISPR/Cas9 guides. (G) Colony counting. Data are the mean ± s.d. (n=3 independent experiments). Student's t-test was used, and two-sided p-values are indicated. (H) Statistical overrepresentation analysis. The Pantherdb tool was used to detect overrepresented GO terms within the genes differentially expressed in control- and Atoh8-KD-derived cell lines. A Fisher's exact two-sided test was used to calculate p-values. (I) Western blot depicting Cdh1 induction during malignant transformation in absence of Atoh8. (J) Picture representing Cresyl-violet immortalized foci at day 30 of foci formation assay starting from control- and Atoh8-KD-derived cell lines. (K) Colony counting. Data are the mean ± s.d. (n=3 independent experiments). Student's t-test was used, and two-sided p-values are indicated. (L) Picture representing Cresyl-violet transformed colonies at day 30 of soft-agar assay starting from cell lines from (K). (M) Colony counting. Data are the mean ± s.d. (n=3 independent experiments). Student's t-test was used, and two-sided p-values are indicated. (N) Picture representing Cresyl-violet immortalized foci at day 30 of foci formation assay starting from cell lines established from 2 independent experiments in control- or Atoh8-KD settings. (O) Colony counting. Data are the mean ± s.d. (n=2 independent experiments).

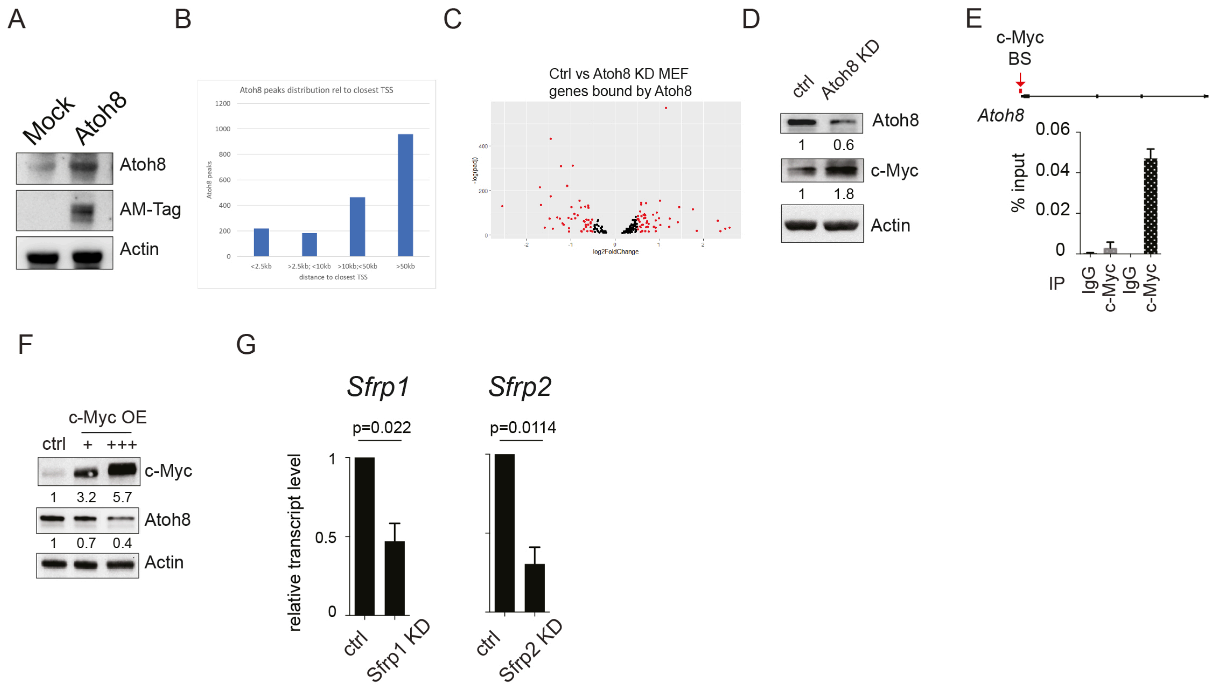


**Supplementary figure 8.** (A) Western blot depicting Atoh8 and AM-Tag detection following the infection of MEF with Mock or Atoh8-AM flagged particles. (B) Screenshot of a ChIP-seq peak, located in the Atoh8 locus itself, in Atoh8 and Mock samples. (C) De novo motif calling with MEME identifies AP-1 DNA-binding motif. (D) Graph depicting the distribution of Atoh8 peaks related to TSS. TSS: Transcription start site. (E) Volcano plot representing the relative expression levels of the 724 genes located near Atoh8 ChIP-seq peaks in Ctrl and Atoh8 KD MEF. (F) Western blot of c-Myc and Atoh8 in Ctrl and Atoh8-KD MEF. (G) Top panel: Scheme depicting c-Myc binding site (BS) on Atoh8 promoter. Bottom panel: q-RTPCR showing levels of DNA immunoprecipitated with control IgG or c-Myc antibody in control or c-Myc-OE MEF. Data, represented as a percentage of Atoh8 DNA levels in ChiP input, are the mean +/- sd of 2 independent experiments. (H) Western blot of c-Myc and Atoh8 in MEF exogenously expressing increasing doses of c-Myc. (I) Q-RTPCR showing *Sfrp1* and *Sfrp2* transcript levels upon downregulation with shRNAs. Data, normalized to control, are the mean +/- sd of 3 independent experiments. Student T-test was used, and two-sided p-values are indicated.

### Table 1: List of sequences

**Genotyping primers**

| Transgene | Primer 1 (5'-3') | Primer 2 (5'-3') | Primer 3 (5'-3') |
|---|---|---|---|
| Col1a1$^{4F2A}$ | CCCTCCATGTGTGACCAAGG | TTGCTCAGCGGTGCTGTCCA | GCACAGCATTGCGGACATG |
| R26$^{rtTA}$ | GCGAAGAGTTTGTCCTCAACC | AAAGTCGCTCTGAGTTGTTAT | GGAGCGGGAGAAATGGATATG |
| LSL-K-ras$^{G12D}$ | CCTTTACAAGCGCACGCAGACTGTAGA | AGCTAGCCACCATGGCTTGAGTAAGTCTGCA | |
| R26-CRE$^{ERT2}$ | TGCCACGACCAAGTGACAGC | CCAGGTTACGGATATAGTTCATG | |
| OCT4-EGFP | CAAGGCAAGGGAGGTAGACA | TGCCAGACAATGGCTATGAG | CCAAAAGACGGCAATATGGT |
| Bcl11b-tdTomato | GCCGGGTACCGAAGACACCAAACCGCTCTT | GCCCGGATCCTCCCCTTGGCAACTACTGAC | |

**shRNA sequences (5'-3')**

| | |
|---|---|
| p53 | CCCACTACAAGTACATGTGTAA |
| FosL1 | CCAGTGCCTTGCATCTCCCTT |
| Atoh8 | CGTCAATTTCACACGTAATTT |
| shSfrp1 | ACTGGCCCGAGATGCTCAAAT |
| shSfrp2 | CGGCATCGAGTACCAGAACAT |

**qPCR primers (5'-3')**

| | Forward primer | Reverse primer |
|---|---|---|
| *Gapdh* | CATGGCCTTCCGTGTTCCTA | GCCTGCTTCACCACCTTCTT |
| *Rplp0* | GCTGATCATCCAGCAGGTGT | GGACACCCTCCAGAAAGCGA |
| *Bcl11b* | GGGAACTCATCACGCCAGAG | TGAGTAGATCAGGGTCGGGG |
| *Atoh8* | CCTCAGCTTCTCCGAGTGTG | CAGGTCACTCCTTCCGTTTCT |
| *Gata4* | TGGAAGACACCCCAATCTCG | TAGTCTGGCAGTTGGCACAG |
| *Sox17* | GACTCCGGTGTGAATCTCCC | TAACACTGCTTCTGGCCTGC |
| *Brachyury (T)* | CGCTGTGACTGCCTACCAGAATG | GAGAGAGAGCGAGCCTCCAAAC |
| *Eomes* | AGCCATGTTTGCCCTAGTCC | GCTTGCTCTCTCCTGAGTCC |
| *Sfrp1* | GGAAGCCTCTAAGCCCCAAG | CATCCTCAGTGCAAACTCGC |
| *Sfrp2* | GTGTCCGAAAGGGACCTGAA | TGACCAGATACGGAGCGTTG |

**Guide CRISPR sequences (5'-3')**

| | Forward sequence |
|---|---|
| Atoh8-1 | GGAAGCACATCCCGGTCCTCG |
| Atoh8-2 | GCCGGGATGTGCTTCATGGCG |
